# A spatiotemporally resolved atlas of mRNA decay in the *C. elegans* embryo reveals differential regulation of mRNA stability across stages and cell types

**DOI:** 10.1101/2024.01.15.575757

**Authors:** Felicia Peng, C Erik Nordgren, John Isaac Murray

## Abstract

During embryonic development, cells undergo dynamic changes in gene expression that are required for appropriate cell fate specification. Although both transcription and mRNA degradation contribute to gene expression dynamics, patterns of mRNA decay are less well-understood. Here we directly measured spatiotemporally resolved mRNA decay rates transcriptome-wide throughout *C. elegans* embryogenesis by transcription inhibition followed by bulk and single-cell RNA-sequencing. This allowed us to calculate mRNA half-lives within specific cell types and developmental stages and identify differentially regulated mRNA decay throughout embryonic development. We identified transcript features that are correlated with mRNA stability and found that mRNA decay rates are associated with distinct peaks in gene expression over time. Moreover, we provide evidence that, on average, mRNA is more stable in the germline compared to in the soma and in later embryonic stages compared to in earlier stages. This work suggests that differential mRNA decay across cell states and time helps to shape developmental gene expression, and it provides a valuable resource for studies of mRNA turnover regulatory mechanisms.

## Introduction

Development is a highly regulated process that relies on the precise spatial and temporal control of gene expression patterns. Although transcriptional regulation is heavily studied, the regulation of mRNA decay can also be complex and necessarily influences mRNA abundance (Alonso 2012). Transcripts are often specified for decay through the binding of RNA and protein factors to cis-regulatory elements within the 3’ untranslated region (UTR) (Ray et al. 2013; Vejnar et al. 2019). Cross-talk between factors can lead to differential mRNA decay, such as RNA-binding proteins stabilizing and protecting transcripts from miRNA-mediated repression (Bhattacharyya et al. 2006; Young et al. 2012). Furthermore, computational analyses predict widespread antagonistic or synergistic actions between RNA-binding proteins and miRNAs in regulating gene expression (Jiang et al. 2013). Translational efficiency and features that influence it, such as codon optimality, are also key determinants of mRNA stability. From yeast to humans, greater translational efficiency is associated with greater transcript stability (Presnyak et al. 2015; Wu et al. 2019). Once transcript turnover is specified, transcripts are ultimately targeted to one of many distinct decay pathways, such as those mediated by the major exoribonucleases XRN1 (5’-3’ decay) and the RNA exosome (3’-5’ decay) (Łabno et al. 2016). Given the potential complexity in the regulation of mRNA degradation, differential mRNA decay in different lineages, cell types, or developmental stages could contribute to distinct gene expression patterns. The degree to which such regulation occurs throughout development is unclear. Measuring how mRNA degradation varies across different cell types and developmental stages would determine the extent of differential mRNA decay during development and provide the basis for identifying mechanisms for this regulation.

The developmental importance of mRNA degradation is underscored by the maternal-to-zygotic transition, in which maternal gene products in the early embryo must be degraded for the control of development to switch to zygotically-encoded products (Vastenhouw et al. 2019). Global studies in both vertebrate and invertebrate model organisms have measured maternal mRNA decay and identified key RNA-binding proteins and small RNAs involved in the clearance of maternal products (Giraldez et al. 2006; Tadros et al. 2007). In addition, studies of zygotic mRNA degradation have uncovered cell differentiation events that are influenced by mRNA decay. For example, timely decay of *glial cells missing* transcripts in the fly embryo is important for appropriate nervous system differentiation (Soustelle et al. 2008), and stabilization of muscle-specific mRNAs by the RNA-binding protein Human antigen R promotes myogenesis (van der Giessen and Gallouzi 2007). The regulation of mRNA degradation for developmental regulators may therefore be important in patterning fates. As studies of zygotic mRNA decay tend to be on a gene-by-gene basis or in the context of cell culture systems, global studies in complex organisms will be necessary to establish the extent of developmentally regulated zygotic decay.

The nematode *Caenorhabditis elegans* provides an ideal system for the global study of zygotic mRNA decay due to its invariant cell lineage, experimental tractability, and conservation of RNA decay machinery with humans. In this study, we provide the first global measurement of mRNA decay rates throughout *C. elegans* embryogenesis at high resolution by pairing a transcription inhibition approach with both bulk and single-cell RNA-sequencing. Our data suggest that the regulation of mRNA decay makes meaningful contributions to specific gene expression patterns across developmental stages and cell types. We further highlight that specific sequence features and RNA-binding proteins may play a role in how this differential mRNA degradation is regulated.

## Results

### Transcription inhibition by actinomycin D allows mRNA decay measurements in C. elegans embryonic cells

To measure zygotic mRNA decay throughout *C. elegans* embryogenesis, we measured changes in gene expression after transcription inhibition (**Figure 1A**). Primary cell cultures of *C. elegans* embryos divide and differentiate in ways that mirror development of intact embryos, with a similar range of cell types differentiating at roughly the same time they would in intact embryos (Edgar 1995; Christensen et al. 2002). We took advantage of this by dissociating mixed-stage embryos into single cells and treating the cells with the transcription inhibitor actinomycin D (actD) in a time course experiment. Transcription inhibition was efficient, as measured by the rapid and dramatic decrease in levels of unspliced transcripts for several housekeeping genes within minutes of actD treatment (**Supplemental Figure S1A**). On the other hand, levels of unspliced transcripts were maintained in cultured cells not treated with actD (**Supplemental Figure S1B**). After an hour of actD treatment, unspliced transcripts remained low and cell viability remained high (> 90%).

**Figure 1.**
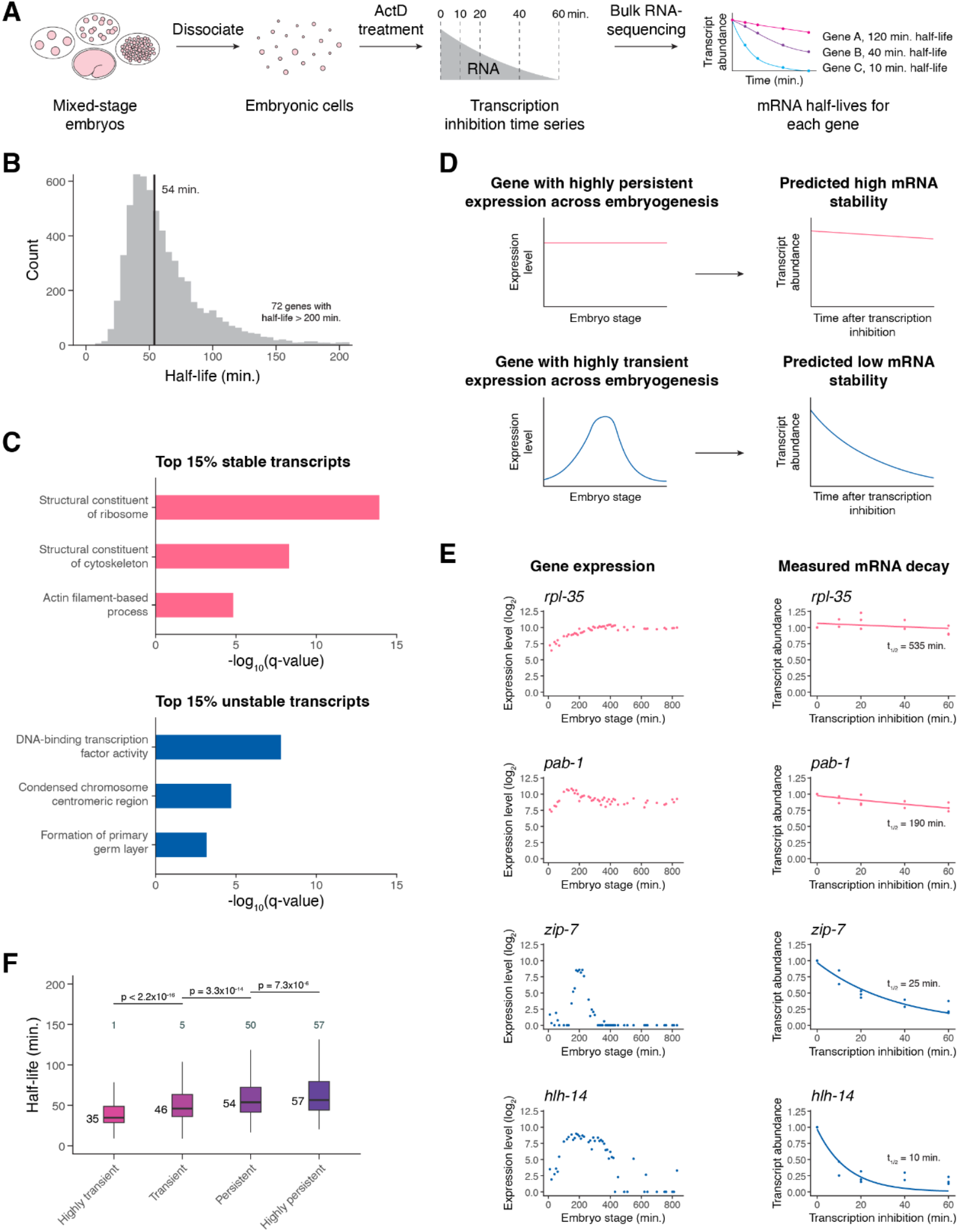
Transcription inhibition by actinomycin D allows mRNA decay measurements in *C. elegans* embryonic cells. (A) A schematic representation of the approach to measure mRNA half-lives in the *C. elegans* embryo using transcription inhibition and bulk RNA-sequencing. (B) Distribution of mRNA half-lives across three biological replicates that met a moderate filtering strategy. Median half-life was 54 minutes (black line). 72 genes with half-lives greater than 200 minutes not shown. (C) Select gene ontology categories enriched among the top 15% stable and unstable transcripts. Background set of genes used was all genes that met our moderate mRNA half-life filtering metric. (D) A schematic representation of a gene with highly persistent expression and a gene with highly transient expression and the predicted overall stability of their transcripts. (E) *Left*. Expression over time of highly persistent (*rpl-35, pab-1*) and highly transient (*zip-7, hlh-14*) genes from a whole-embryo RNA-sequencing time series (Hashimshony et al. 2015). *Right*. The measured mRNA decay of the corresponding genes from our bulk RNA-sequencing data, with each point representing normalized transcript abundance from one of three biological replicates. (F) Box plots showing the mRNA half-life distributions of genes characterized as highly transient, transient, persistent, or highly persistent transcriptome-wide. Numbers to the left of the box plots are median half-lives within each group. Numbers above the box plots are the number of genes with half-lives greater than 150 minutes within each group. P-values comparing median half-lives were calculated using the Wilcoxon rank sum test.

To determine half-lives transcriptome-wide, we measured mRNA levels in cells after 0, 10, 20, 40, and 60 minutes of actD treatment. After each time point, samples were spiked with External RNA Controls Consortium (ERCC) control transcripts (Baker et al. 2005) before RNA was isolated and analyzed using total RNA-sequencing after ribosomal RNA depletion (**Supplemental Table S1**). An mRNA half-life was calculated for each gene by fitting ERCC-normalized gene expression over time to an exponential decay equation across all replicates. The measured half-lives had a median of 54 minutes and varied by more than an order of magnitude from gene to gene, with most ranging from ∼20 to 200 minutes (**Figure 1B, Supplemental Table S2**).

Several pieces of evidence indicate that these half-lives are reproducible and biologically meaningful. First, calculating half-lives separately for each of three biological replicates yielded reproducible half-life estimates (Spearman’s ρ > 0.7 for all pairwise comparisons of well-measured genes) (**Supplemental Figure S1C-S1E**). We focused our downstream analyses on high confidence half-life measurements by implementing a filtering strategy based on an initial count of 30 for each gene and coefficient of variation and 95% confidence interval thresholds for half-lives across biological replicates (**Supplemental Figure S1F, see Methods**).

Second, genes with the slowest or fastest mRNA decay rates were enriched for different functional categories. Gene ontology analysis found an enrichment among the most stable transcripts for those encoding proteins with housekeeping functions, such as ribosomal and cytoskeletal proteins (**Figure 1C, Supplemental Figure S2A**). In contrast, among the most unstable mRNAs there was an enrichment for those encoding proteins involved in dynamic developmental processes, such as transcription factor activity and the cell cycle. (**Figure 1C, Supplemental Figure S2B**). These functional associations with RNA stability generally agree with those found in previous studies of decay in other systems (Yang et al. 2003; Narsai et al. 2007; Thomsen et al. 2010; Burow et al. 2015).

Third, we found that RNA stability correlates with temporal dynamics in untreated embryonic cells. High mRNA stability would be one way in which a gene could maintain persistent expression over time, while rapid drops in expression for transiently expressed genes would necessarily be facilitated by mRNA degradation (**Figure 1D**). Thus, we tested whether transiently expressed genes have shorter mRNA half-lives compared to persistently expressed genes. We defined genes as having either transient or persistent expression over time from a previously published *C. elegans* embryo single cell atlas (Packer et al. 2019) based on the magnitude and frequency of expression decrease from mother to daughter cell states (see **Methods**). For example, transcription factor genes *zip-7* and *hlh-14* were characterized as having highly transient expression due to the loss of expression from parent to daughter cell in multiple lineages (**Supplemental Figure S2C, S2D**). On the other hand, the genes *rpl-35* and *pab-1* were characterized as having highly persistent expression due to the maintenance of expression from parent to daughter cell across multiple lineages (**Supplemental Figure S2E, S2F**). These genes show similar temporal dynamics in a separate *C. elegans* whole embryo time series dataset (Hashimshony et al. 2015); *zip-7* and *hlh-14* increase to peaks in expression before dropping off rapidly, while *rpl-35* and *pab-1* have fairly constant expression over time (**Figure 1E**). In agreement with our expectations, we found that *zip-7* and *hlh-14* transcripts decayed much more rapidly than those of *rpl-35* and *pab-1* (**Figure 1E**). Across the transcriptome, highly transient genes generally had the shortest mRNA half-lives while highly persistent genes generally had the longest mRNA half-lives (**Figure 1F**). These findings are consistent with mRNA decay having an important contribution to gene expression dynamics.

Overall, the results from our bulk RNA-sequencing data suggest that our transcription inhibition approach provides biologically meaningful and reproducible half-lives for zygotic transcripts in the *C. elegans* embryo.

### Transcript stability correlates with specific sequence features

To identify features that differ between stable and unstable transcripts, we tested for correlations between various transcript attributes and our mRNA stability measurements. We identified numerous features associated with transcript stability.

Codon optimality concerns the nonuniform decoding rate of codons by the ribosome, with a codon being considered optimal when its cognate tRNA is prevalent and allows for rapid decoding (Bae and Coller 2022). In examining codon optimality scores between the top 15% most stable and unstable transcripts, we found that the most stable transcripts had significantly greater codon optimality than the least stable transcripts (**Figure 2A**).

**Figure 2.**
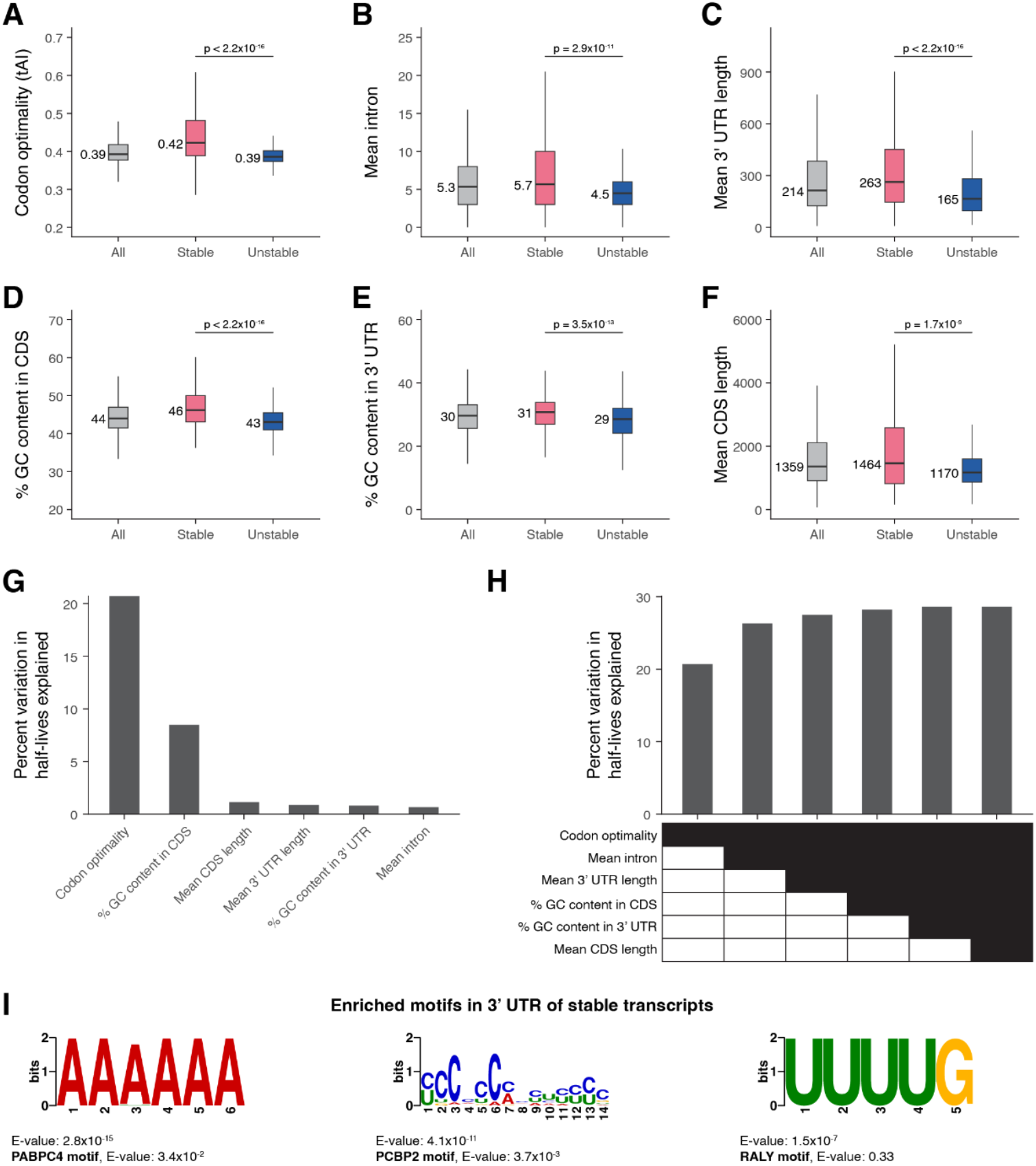
Transcript stability correlates with specific sequence features. The relationship between transcript stability and different sequence features was examined using the top 15% stable and unstable transcripts identified in our bulk RNA-sequencing data. The box plots compare different features between all, stable, and unstable transcripts. Numbers to the left of the box plots are median values within each group. P-values comparing median values were calculated using the Wilcoxon rank sum test. Outliers not shown. (A) Distribution of codon optimality scores, as measured using tRNA adaptation index (tAI) values for each gene. Higher tAI value corresponds to greater codon optimality. (B) Distribution of the number of introns averaged across all splice isoforms of a gene. (C) Distribution of 3’ UTR length averaged across all 3’ UTR isoforms of a gene. (D) Distribution of percent GC content in the coding sequence of genes. For genes with multiple coding sequence isoforms, the longest isoform was used. (E) Distribution of percent GC content in the 3’ UTR of genes. For genes with multiple 3’ UTR isoforms, the longest isoform was used. (F) Distribution of coding sequence length averaged across all splice isoforms of a gene. (G) Linear regression was used to identify the percent of variation in mRNA half-lives explained by individual sequence features. (H) Multiple linear regression was used to identify the percent of variation in mRNA half-lives explained by combinations of different sequence features. (I) The *de novo* motif-finding program MEME (Bailey et al. 2015) identified three motifs enriched in the 3’ UTRs of the top 15% stable transcripts compared to the 3’ UTRs of the top 15% unstable transcripts. For genes with multiple 3’ UTR isoforms, the longest isoform was used. The best match of each *de novo* motif to known motifs in mammals (Ray et al. 2013) is noted.

We next examined whether intron number is correlated with mRNA stability, since the presence of introns has been associated with higher translation (Shaul 2017). We found that unstable mRNAs had significantly fewer introns compared to stable mRNAs, with a median of 4.5 and 5.7 mean number of introns per gene, respectively (**Figure 2B**). This was correlated with differences in overall gene length; mean coding sequence length per gene was correlated with mRNA stability and significantly shorter for unstable transcripts (**Figure 2F**).

Since 3’ UTRs can influence mRNA stability (Mayr 2019), we tested for 3’ UTR characteristics correlated with half-lives. Unstable transcripts had significantly shorter 3’ UTR lengths compared to stable transcripts, with a median difference of 98 base pairs (**Figure 2C**). Nucleotide composition of both the 3’ UTR and coding sequence also varied with stability, with higher GC content for stable transcripts (**Figure 2D, 2E**).

We modeled whether these different transcript features provide independent information about mRNA half-lives by using linear regression. Examining the contribution of individual features showed that codon optimality explained the largest fraction of variation in mRNA stability (∼21%) (**Figure 2G**). Other features such as mean number of introns, mean 3’ UTR length, and GC content in the coding sequence provided independent information about variation in mRNA stability (**Figure 2H**). Most of the variation in mRNA stability, however, remains unexplained by the examined sequence features. To identify sequence motifs that may contribute to differences in transcript stability, we used the *de novo* motif-finding program MEME (Bailey et al. 2015) to identify motifs differentially enriched in the 3’ UTRs of the most stable and unstable transcripts. No statistically significant motifs were enriched for unstable transcripts, but several motifs were found to be differentially enriched for stable transcripts (**Figure 2I, Supplemental Figure S3A**). Comparing these motifs against a database of known motifs (Ray et al. 2013) identified potential regulators of RNA stability (**Figure 2I, Supplemental Figure S3B**). The YCCCCCMHYUYYYC motif resembles poly(C)-binding protein (PCBP) family binding sites, the AAAAAA motif resembles poly(A)-binding protein (PABP) family binding sites, and the UUUUG motif resembles heterogeneous nuclear ribonucleoprotein (hnRNP) family binding sites. Notably, RNA-binding proteins within these families have previously been shown to have transcript-stabilizing effects in other systems (Makeyev and Liebhaber 2002; Geuens et al. 2016; Passmore and Coller 2021).

These findings highlight that stable and unstable transcripts have several distinct sequence features, suggesting that multiple mechanisms may influence their differences in stability. Codon optimality appears to be the single strongest predictor of mRNA stability, in line with translational efficiency being a key determinant of mRNA degradation. Additional complexity to the regulation of mRNA decay may result from other binding sites for regulatory RNAs and proteins that act in specific contexts.

### High transcript accumulation is associated with increased mRNA stability

Throughout *C. elegans* embryogenesis, many genes accumulate high transcript levels. While this requires high transcription rates, we asked whether rapid transcript accumulation is also associated with increased mRNA half-life, as such accumulation would be difficult to achieve in the presence of rapid RNA turnover. Genes with lower accumulation rates, on the other hand, could be more tolerant of RNA turnover rates (**Figure 3A**). For example, moderate transcription rate with moderate mRNA half-life or high transcription rate with high RNA turnover could yield similar slopes of RNA versus time.

**Figure 3.**
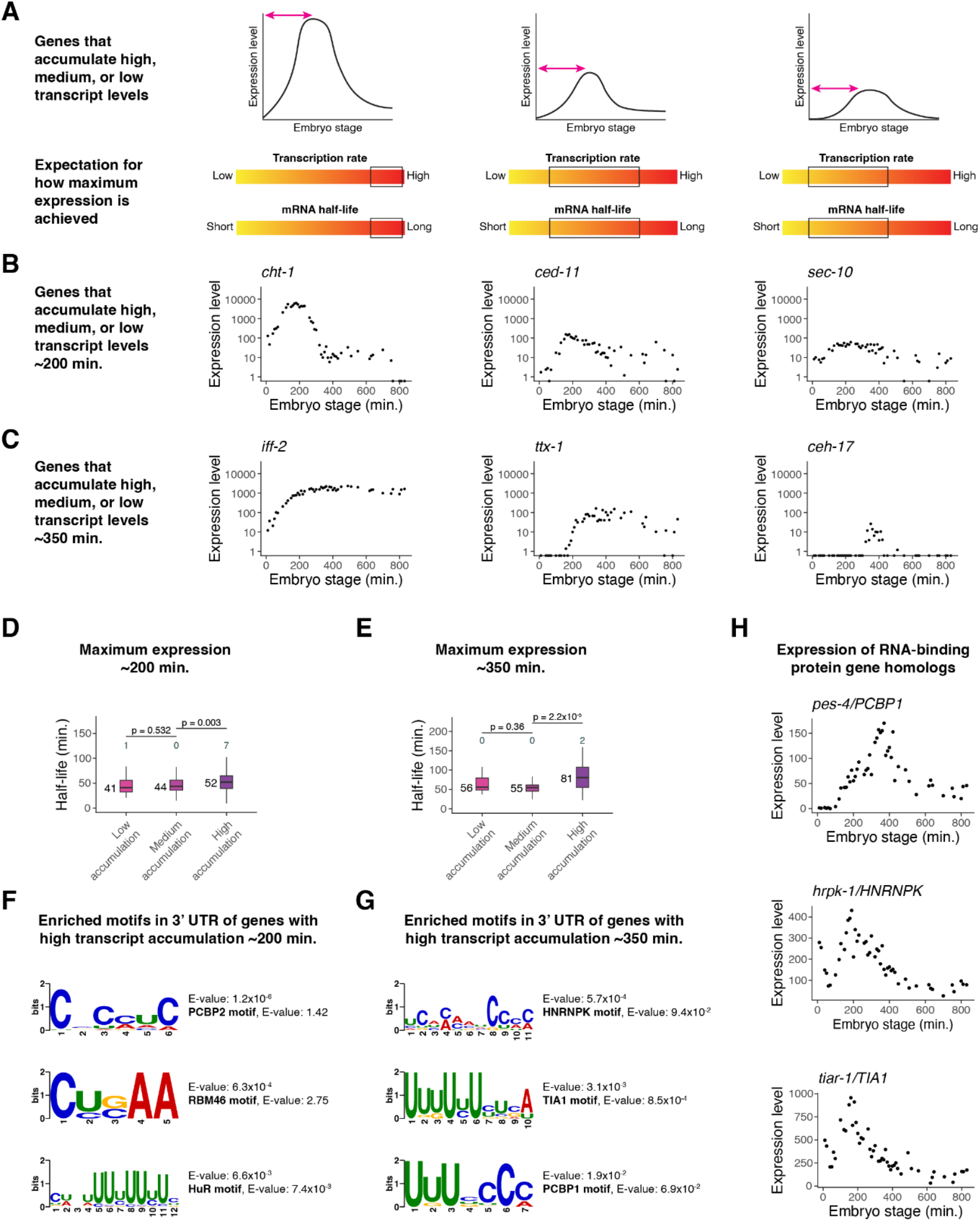
High transcript accumulation is associated with increased mRNA stability. (A) A schematic representation of genes that accumulate high, medium, or low transcript levels and the expected range of transcription and mRNA decay rates that could contribute to such accumulation. (B) Examples of genes that peak in expression ∼200 minutes after the four-cell stage with high, medium, and low transcript accumulation, from left to right. Expression data taken from a whole-embryo RNA-sequencing time series (Hashimshony et al. 2015). (C) Examples of genes that peak in expression ∼350 minutes after the four-cell stage with high, medium, and low transcript accumulation, from left to right. Expression data taken from a whole-embryo RNA-sequencing time series (Hashimshony et al. 2015). (D) Box plots showing the mRNA half-life distributions of genes that peak in expression ∼200 minutes after the four-cell stage to low, medium, and high transcript levels. (E) Box plots showing the mRNA half-life distributions of genes that peak in expression ∼350 minutes after the four-cell stage with low, medium, and high transcript levels. (F) The *de novo* motif-finding program MEME (Bailey et al. 2015) identified three motifs enriched in the 3’ UTRs of genes with high transcript accumulation compared to the 3’ UTRs of genes with low transcript accumulation ∼200 minutes after the four-cell stage. For genes with multiple 3’ UTR isoforms, the longest isoform was used. The best match of each *de novo* motif to known motifs in mammals (Ray et al. 2013) is noted. (G) The *de novo* motif-finding program MEME (Bailey et al. 2015) identified three motifs enriched in the 3’ UTRs of genes with high transcript accumulation compared to the 3’ UTRs of genes with low transcript accumulation ∼350 minutes after the four-cell stage. For genes with multiple 3’ UTR isoforms, the longest isoform was used. The best match of each *de novo* motif to known motifs in mammals (Ray et al. 2013) is noted. (H) The gene expression patterns for three putative RNA-binding protein genes in *C. elegans* from a whole-embryo RNA-sequencing time series (Hashimshony et al. 2015). The genes are homologs of mammalian RNA-binding protein genes whose corresponding proteins are known to bind motifs similar to those discovered in (F) and (G). Numbers to the left of the box plots are median half-lives within each group. Numbers above the box plots are the number of genes with half-lives greater than 125 minutes (D) or 175 minutes (E) within each group. P-values comparing median half-lives were calculated using the Wilcoxon rank sum test.

Using staged embryo RNA-sequencing data <u>(Hashimshony et al. 2015)</u>, we identified 985 dynamic genes whose expression peaks at ∼200 minutes past the four-cell stage. We categorized these genes as having high (top 20%), medium (middle 20%), or low (bottom 20%) transcript accumulation rates (**Figure 3B**). For example, the high-accumulation gene *cht-1* peaks at an expression level of around 6,300 transcripts per million (TPM), which is in the top 1% of maximum expression across all genes and accounts for ∼0.6% of transcripts in the embryo. On the other hand, the medium-accumulation gene *ced-11* peaks at around 150 TPM, and the low-accumulation gene *sec-10* peaks at around 60 TPM.

We found that genes with high accumulation had a median mRNA half-life about 27% longer than that of genes with low accumulation (median t_1/2_ of 52 vs 41 minutes; **Figure 3D**). Similar results were observed for genes whose expression peaks later, around 350 minutes past the four-cell stage (**Figure 3C**). At this stage, high-accumulation genes had a median mRNA half-life about 45% longer than that of low-accumulation genes (median t_1/2_ of 81 vs 56 minutes) (**Figure 3E**).

We identified motifs differentially enriched in the 3’ UTRs of high-accumulation genes compared to the 3’ UTRs of low-accumulation genes using the *de novo* motif finding program MEME (Bailey et al. 2015). Searching an existing motif database (Ray et al. 2013) identified possible binding factors for each of these motifs (**Figure 3F, 3G; Supplemental Figure S4A, S4C**). High-accumulation genes peaking at ∼200 minutes were enriched for potential embryonic lethal abnormal vision-like (ELAVL) and RNA-binding motif (RBM) family binding sites, those peaking at ∼350 minutes were enriched for potential hnRNP and T-cell intracellular antigen 1 (TIA1) family binding sites, and high-accumulation genes at both stages were enriched for likely PCBP binding sites (**Supplemental Figure S4B, S4D**). Several of the corresponding RNA-binding protein genes have *C. elegans* homologs that have distinct peaks in mRNA expression throughout embryogenesis (**Figure 3H**).

Overall, dynamically expressed genes with high transcript accumulation rates tend to have longer mRNA half-lives. This suggests that regulation of both transcription and mRNA decay help these genes accumulate to high transcript levels.

### Single-cell RNA-sequencing allows measurement of mRNA half-lives at high resolution throughout C. elegans embryogenesis

While our bulk RNA-sequencing data supports the idea that many zygotic genes vary in their decay rates, it does not allow us to identify changes in transcript stability for the same gene in different cell types or developmental stages. To globally identify developmentally regulated mRNA decay in the *C. elegans* embryo, we combined actD transcription inhibition with single-cell RNA-sequencing (**Figure 4A**).

**Figure 4.**
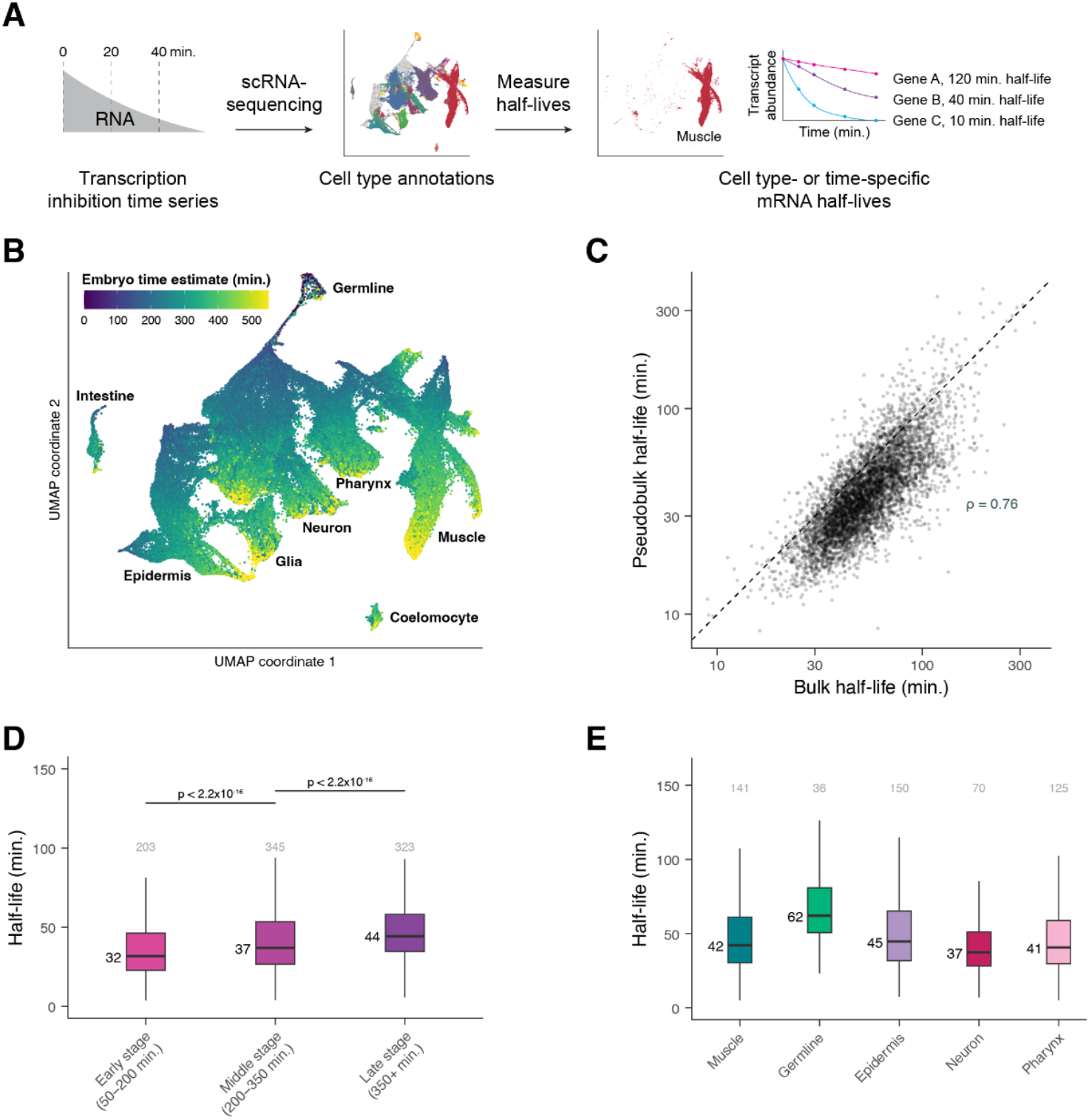
Single-cell RNA-sequencing allows measurement of mRNA half-lives at high resolution throughout *C. elegans* embryogenesis. (A) A schematic representation of the approach to measure mRNA half-lives in the *C. elegans* embryo using transcription inhibition and single-cell RNA-sequencing. (B) UMAP projection of the integrated dataset of three biological replicates. Cells are colored by the age of the embryo from which a cell was produced, estimated from correlations to a whole-embryo RNA-sequencing time series (Hashimshony et al. 2015). Trajectories corresponding to major cell types are labeled. (C) Scatter plot comparing the mRNA half-lives calculated in pseudobulk from the single-cell data and the half-lives calculated from the bulk-cell data on a log-log scale. Spearman correlation coefficient = 0.76. Dashed line is the x = y line. (D) Box plots showing the stage-specific mRNA half-life distributions of genes within Early-, Middle-, and Late-stage cells (50-200, 200-350, and 350+ minutes after the four-cell embryo stage, respectively). (E) Box plots showing the cell type-specific mRNA half-life distributions of genes within muscle, germline, epidermis, neuron, and pharynx. Numbers above the box plots are the number of genes with half-lives greater than 100 minutes (D) or 150 minutes (E) within each group. P-values comparing median half-lives were calculated using the Wilcoxon rank sum test.

We dissociated mixed-stage populations of *C. elegans* embryos into single cells and cultured them for a time course in the presence or absence of actD, as was done for the bulk data described above. mRNA levels were then measured in single cells using the 10x Genomics single-cell RNA-sequencing platform. Cells from 0-minute (untreated), 20-minute actD-treated, and 40-minute actD-treated cells were used to calculate mRNA half-lives. An additional 40-minute untreated time point served as a control for the impact of cell culturing. We used this slightly shorter time course to allow for more accurate annotation of actD-treated cells, as modeling from the bulk data indicated that these reduced time points were sufficient to accurately estimate mRNA half-lives for most genes (**Supplemental Figure S5D**). We performed the entire experiment across three biological replicates and in total captured 183,777 cells (**Supplemental Table S1**).

We integrated the three biological replicates using Seurat, and both manual and automated annotation approaches were used to identify each cell’s terminal cell type or lineage identity and developmental stage (in minutes from the 4-cell stage) (**Supplemental Figure S6A**, see **Methods**). We projected the integrated dataset to two dimensions using the Uniform Manifold Approximation and Projection (UMAP) algorithm (Becht et al. 2019; McInnes et al. 2020), which organized the cells by type and stage (**Figure 4B**). From a central group of progenitor cells, trajectories formed that generally reflected progression through embryogenesis and that branched out into major cell types. Projecting data from individual biological replicates using the UMAP algorithm revealed similar cell clustering (**Supplemental Figure S5A-S5C**). Importantly, transcription inhibition did not appear to significantly affect the fraction of cells annotated as each major cell type across biological replicates (**Supplemental Figure S6B**).

Half-lives were calculated in a manner similar to those calculated for our bulk data. However, as we were not able to use ERCC control transcripts with the single-cell RNA-sequencing approach, expression of each gene was normalized to the expression of ribosomal protein gene transcripts. Ribosomal protein genes were chosen since their transcripts were highly stable in our spike-in control-normalized bulk data (median t_1/2_ = 295.5 minutes). We then corrected for the expected decay of transcripts encoding ribosomal proteins in our bulk data (**Supplemental Figure S5E**, see **Methods**). Using this normalization approach on the bulk data resulted in half-lives that were highly correlated to those calculated by ERCC normalization (Pearson’s r = 0.999, **Supplemental Figure S5F, Supplemental Table S3**), and applying the same approach to our single-cell data prevented an artificial inflation of half-lives for slow-decaying transcripts. Final pseudobulk half-lives for well-measured transcripts from each of three single-cell biological replicates were well-correlated with one another (Spearman’s ρ > 0.7 for all pairwise comparisons) (**Supplemental Figure S5G-S5I**) and with half-lives calculated from the bulk data (Spearman’s ρ = 0.76, **Figure 4C**). This indicates that the single-cell RNA-sequencing approach captures comparable half-lives to those determined from the bulk approach.

To examine how mRNA half-life changes across development, we focused first on broad developmental stages and cell types. Half-lives for these cell subsets were calculated in pseudobulk as before. To examine whether mRNA half-lives are regulated across developmental stages, we divided embryogenesis into three major phases using estimated embryo time: Early (50-200 minutes), Middle (200-350 minutes), and Late (350+ minutes). Early and Middle stages correspond to embryonic cleavage and mostly contain progenitor cell states, while Late stage contains mostly terminally differentiating cells. Overall, we observed increased mRNA stability over time; median half-life in Late cells was about 38% higher than in Early cells, with Middle cells having an intermediate range of half-lives (**Figure 4D, Supplemental Table S4-6**). Similarly, we compared half-lives in broad cell classes (muscle, germline, epidermis, neuron, and pharynx) (**Supplemental Table S7-12**). The most dramatic difference observed was the overall longer half-lives in the germline compared to in somatic cell types (**Figure 4E, Supplemental Figure S6C**). Half-life distributions were significantly different between somatic cell types as well, but the magnitude of these differences was generally smaller (**Figure 4E, Supplemental Figure S6C**). Together these observations suggest that global mRNA turnover rates are regulated across both developmental stages and cell types.

### Differential mRNA decay occurs throughout different developmental stages of C. elegans embryogenesis

While transcripts on average increased in stability over time (**Figure 4D**), we identified a subset of transcripts that decayed faster at later stages of embryogenesis (**Figure 5A**). Genes with decreasing mRNA stability over time were enriched for terms such as structural constituent of chromatin, transporter activity, and cilium organization (**Figure 5B, Supplemental Figure S7A-S7C**).

**Figure 5.**
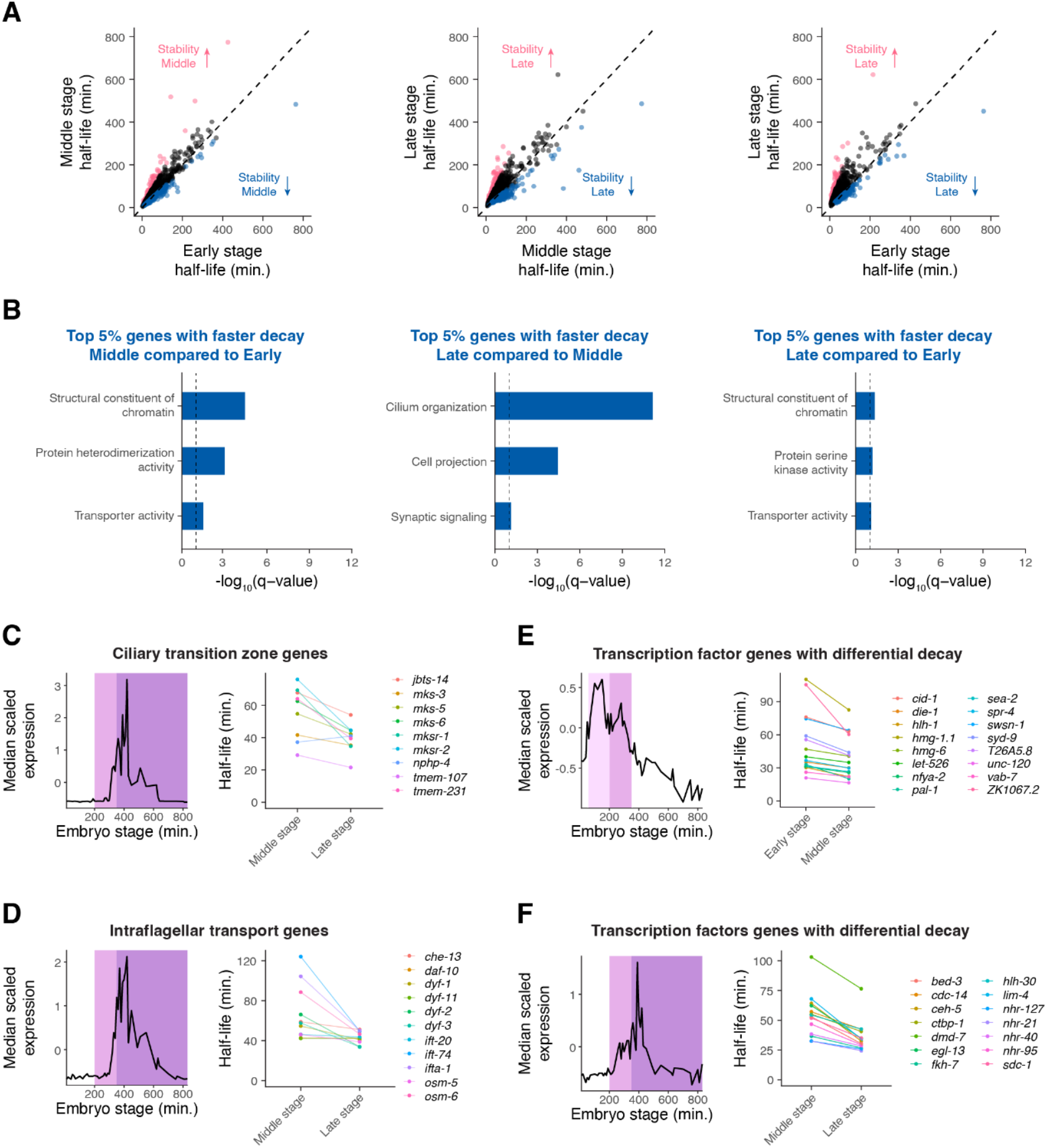
Differential mRNA decay occurs throughout different developmental stages of *C. elegans* embryogenesis. (A) Scatter plots comparing mRNA half-lives specific to Early-, Middle-, and Late-stage cells in all pairwise comparisons. Each point represents a gene. Blue points correspond to the top 5% of genes with faster mRNA decay in the later stage compared to in the earlier stage. Pink points correspond to the top 5% of genes with slower mRNA decay in the later stage compared to in the earlier stage. Dashed line is the x = y line. (B) Select gene ontology categories enriched among the top 5% of genes with faster mRNA decay in the later stage of embryogenesis compared to the earlier stage of embryogenesis. Background set of genes used in each comparison was shared genes we were able to calculate stage-specific mRNA half-lives for between the relevant stages. (C, D, E, F) *Left*. Median scaled expression of gene subset using data from a whole-embryo RNA-sequencing time series (Hashimshony et al. 2015). Pink shading spans the Early stage, light purple shading spans the Middle stage, and dark purple shading spans the late stage. *Right*. Plot displaying the change in mRNA half-lives from earlier to later stage for gene subset.

To illustrate this differential mRNA decay, we examined the dynamics of 70 previously identified core cilia component genes (Brocal-Ruiz et al. 2023), since genes annotated with ciliary functions were enriched among genes with faster mRNA decay over time. These genes generally peaked in expression around 400 minutes, around the end of our Middle stage (**Supplemental Figure S8A**). This corresponds well to the time when many sensory neurons generate their cilia (Nechipurenko and Sengupta 2017). While the average transcript becomes more stable in Late-vs Middle-stage cells, the average cilia transcript becomes less stable or maintains a similar half-life during this period (**Supplemental Figure S8A**). This pattern was especially pronounced for transcripts encoding ciliary transition zone (Li et al. 2015; Roberson et al. 2015; Lambacher et al. 2016; Li et al. 2016) and intraflagellar transport proteins (Qin et al. 2001; Haycraft et al. 2003; Blacque et al. 2006; Efimenko et al. 2006; Burghoorn et al. 2007; Kobayashi et al. 2007; Ou et al. 2007; Kunitomo and Iino 2008; Mijalkovic et al. 2018; De-Castro et al.) (**Figure 5C, 5D**). The ciliary transition zone is a domain at the base of cilia that controls the entry and exit of ciliary proteins needed for signal transduction (Li et al. 2016). On the other hand, intraflagellar transport mediates protein trafficking along a microtubular axoneme for the assembly and maintenance of cilia (Wang et al. 2021). Our observations suggest that these cilia components are produced at high levels during ciliogenesis, with transcripts destabilized once they are no longer needed at high levels.

Many putative transcription factor genes (Fuxman Bass et al. 2016) displayed changes in mRNA decay over time. We identified 16 transcription factor transcripts with decreasing stability in Middle-stage cells and 14 with decreasing stability in Late-stage cells compared with the prior stage. On average these genes show decreasing expression during the time when their transcript stability is lower (**Figure 5E-5F**). Thus, differential mRNA decay may allow transcripts of developmental regulators to degrade in a timely manner after accumulating to a certain threshold.

As the median scaled expression patterns of these ciliary and transcription factor genes have distinct peaks, this suggests that differential mRNA degradation may contribute to specific patterns of expression throughout embryogenesis. Indeed we found that similar analyses to all zygotic-only expressed genes with faster mRNA decay over time identified peaks in expression corresponding well to the time when stability decreases (**Supplemental Figure S8B**). Importantly, the expression patterns of genes with increased mRNA turnover over time were distinct from those observed for genes with decreased mRNA turnover over time and did not peak as strongly (**Supplemental Figure S8C**).

These results highlight that differential mRNA decay over time is correlated with dynamic gene expression patterns throughout embryogenesis. This suggests that the regulation of mRNA degradation makes important contributions to developmental gene expression.

### mRNA degradation may contribute to transcription factor dynamics at both the RNA and protein levels

Because transcription factors drive developmental gene expression, their mRNA dynamics are of particular interest. Across all developmental stages, transcripts encoding transcription factors (Fuxman Bass et al. 2016) decay more rapidly compared to those encoding other genes (**Figure 6A**). This suggests that transcripts encoding transcription factors may need to achieve faster turnover on average throughout embryogenesis.

**Figure 6.**
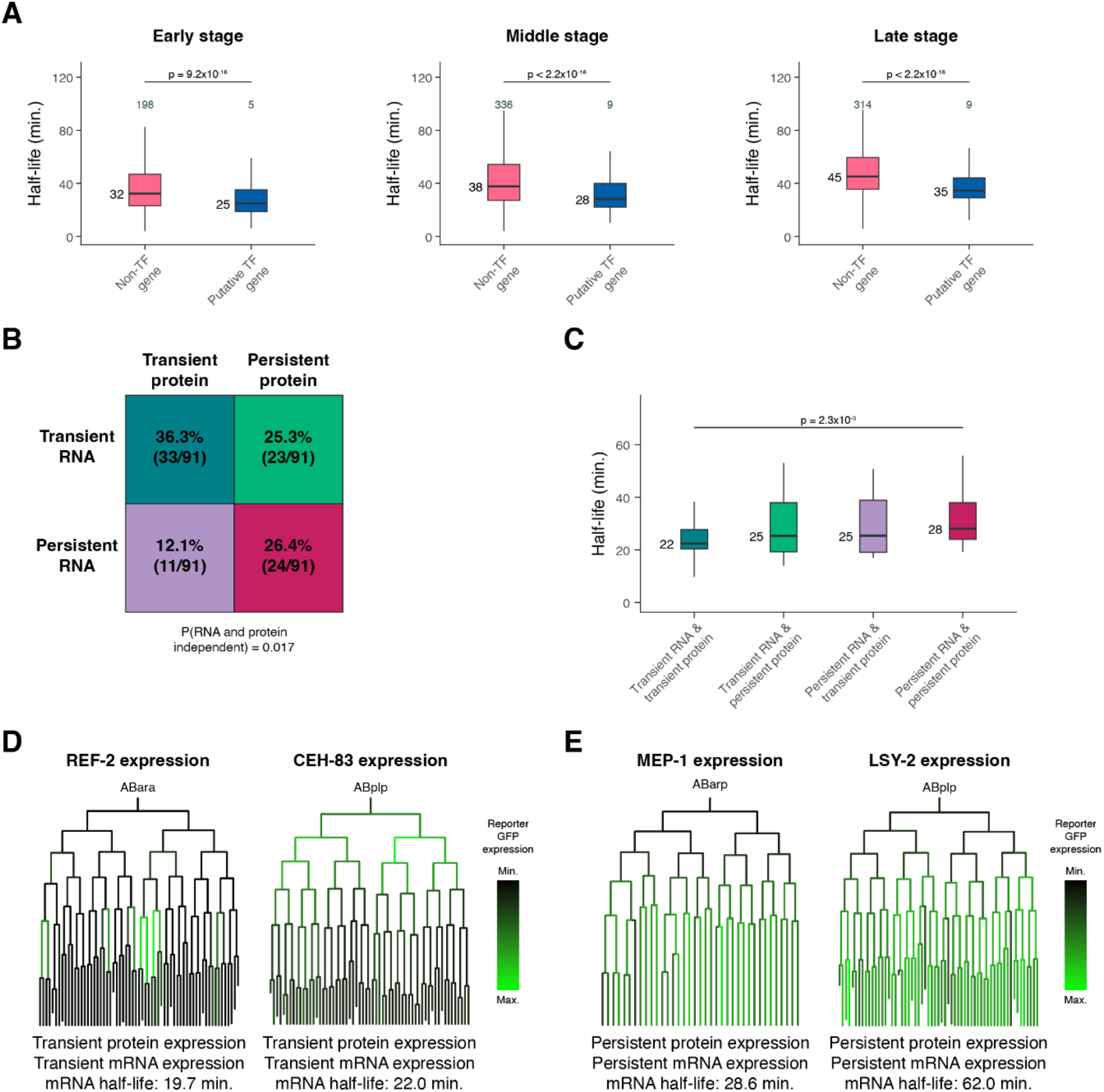
mRNA degradation may contribute to transcription factor dynamics at both the RNA and protein levels. (A) Box plots showing the mRNA half-life distributions of transcription factor genes and all other genes in Early-, Middle-, and Late-stage cells. (B) Grid displaying the percentage of transcription factors with transient or persistent mRNA expression and transient or persistent protein expression. The probability that RNA and protein dynamics are independent of one another was calculated using Fisher’s exact test. (C) Box plots showing the single-cell RNA-sequencing pseudobulk mRNA half-life distributions of transcription factors with transient mRNA and protein expression, transient mRNA and persistent protein expression, persistent mRNA and transient protein expression, and persistent mRNA and protein expression. (D, E) Sublineages with coloring representing reporter GFP expression from a single-cell transcription factor protein expression atlas of the *C. elegans* embryo (Ma et al. 2021). Protein and mRNA dynamics are characterized below, along with pseudobulk measured mRNA half-life. Numbers to the left of the box plots are median half-lives within each group. Numbers above the box plots in (A) are the number of genes with half-lives greater than 100 minutes within each group. P-values comparing median half-lives were calculated using the Wilcoxon rank sum test.

To test whether mRNA decay dynamics are associated with protein expression over time, we took advantage of an existing single-cell transcription factor protein expression atlas of the *C. elegans* embryo (Ma et al. 2021). This study used live-imaging and automated cell lineage tracing of strains expressing almost 300 transcription factor-GFP fusion proteins to track protein expression at single-cell and ∼1-minute temporal resolution. We categorized transcription factor proteins as either transiently or persistently expressed based on the occurrence of loss or maintenance of expression from parent to daughter cells. Similarly, we categorized transcription factors as transiently or persistently expressed at the mRNA level using our single-cell RNA-sequencing atlas (Packer et al. 2019). Overall, among the 91 transcription factors present in both protein and mRNA datasets, there was a significant enrichment for concordant protein and RNA dynamics: 36.3% had both transient protein and mRNA expression, and 26.4% had both persistent protein and mRNA expression (**Figure 6B**, P = 0.017). A smaller number had discordant dynamics, including 12.1% with transient protein but persistent mRNA, and 25.3% with persistent protein but transient mRNA (**Supplemental Table S13**). These discordant cases might reflect interesting classes of post-transcriptional regulation.

We hypothesized that transcription factors with concordant RNA and protein dynamics might reflect stronger regulation by RNA degradation. Consistent with this, transcription factors with transient protein and mRNA had shorter mRNA half-lives on average compared to transcription factors with persistent protein and mRNA (**Figure 6C**). This suggests that transcript turnover may contribute to protein dynamics by influencing overall mRNA expression dynamics for these genes. For example, the transcription factors REF-2 and CEH-83 were both transiently expressed, with protein expression being lost across many lineages (**Figure 6D**). The corresponding mRNAs were also transiently expressed (**Supplemental Figure S9A-S9B**), with mRNA half-lives of 19.7 and 22.0 minutes, respectively. The proteins MEP-1 and LSY-2, on the other hand, were both persistently expressed (**Figure 6E**). The corresponding mRNAs were also persistently expressed (**Supplemental Figure S9C-S9D)**, with half-lives of 28.6 and 62.0 minutes, respectively.

These results suggest that mRNA degradation may contribute to the protein dynamics of developmental regulators. In many cases, the regulation of transcription and mRNA decay appear to act in concert to have a predictable impact on corresponding protein levels. However, it is not uncommon for RNA dynamics to differ from protein dynamics, highlighting the complexity associated with the regulation of protein expression.

### mRNA stability is correlated with cell type-specific functions

As described above, different somatic cell types have more similar mRNA half-life distributions to one another than they do to germline cells (**Figure 4E**). We next asked what categories of genes have faster or slower mRNA decay within each cell class. We identified cell type-specific genes within each class and tested whether their stability differed from broadly expressed genes (see **Methods**). For muscle cells and neurons, cell type-specific transcripts had significantly longer half-lives compared to broadly expressed transcripts, while for the other somatic cell classes, specific and broadly expressed genes had similar stability distributions (**Figure 7A-7B, Supplemental Figure S10A**).

**Figure 7.**
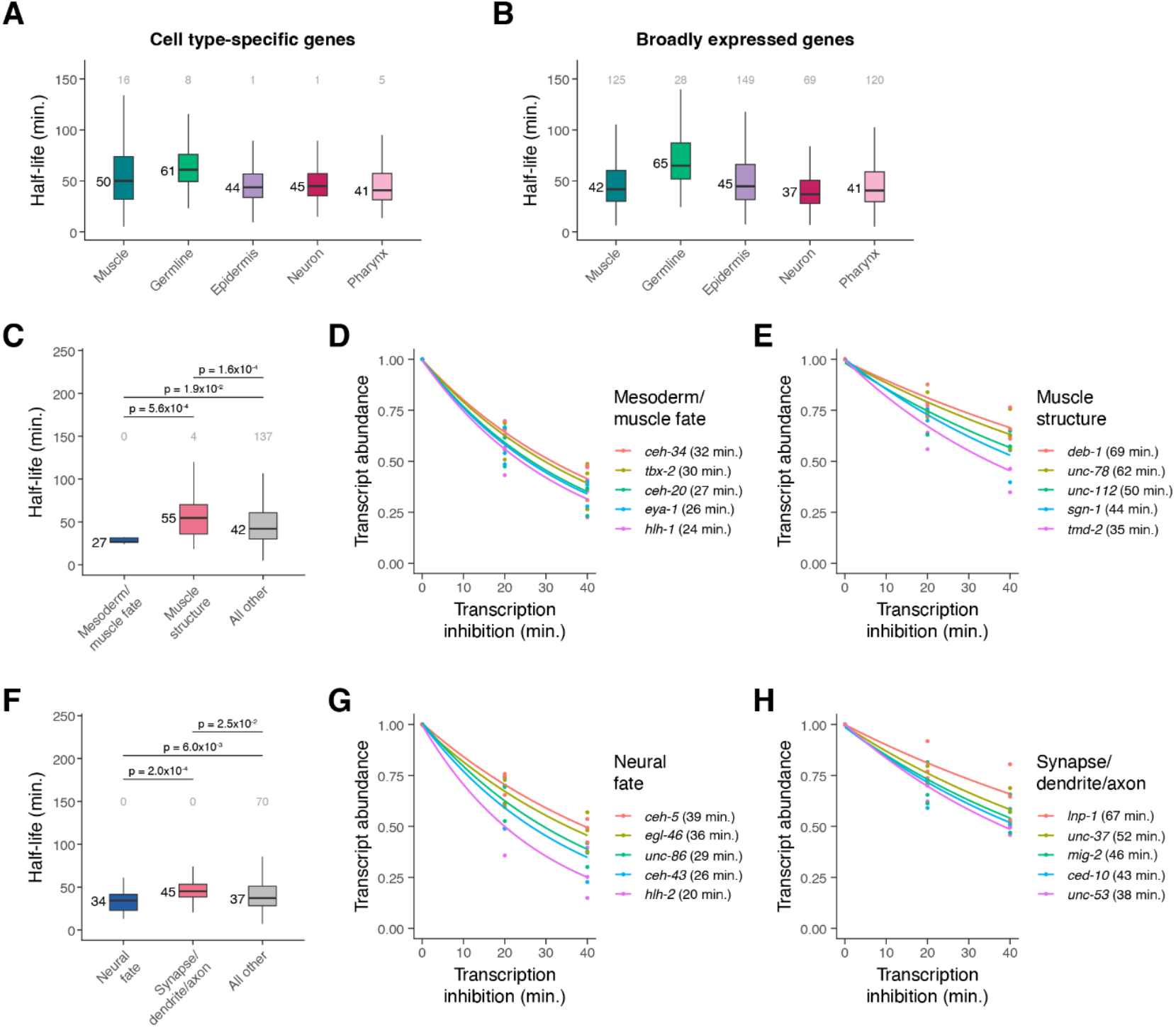
mRNA stability is correlated with cell type-specific functions. (A) Box plots showing the mRNA half-life distributions of cell type-specific genes within muscle, germline, epidermis, neuron, and pharynx cells. (B) Box plots showing the mRNA half-life distributions of broadly expressed genes within muscle, germline, epidermis, neuron, and pharynx. (C) Box plots showing the muscle-specific mRNA half-life distributions of mesoderm/muscle fate-specifying transcription factor genes, muscle structure genes, and all other genes. (D, E) Scatter plots of the normalized transcript abundance of the mesoderm/muscle fate-specifying transcription factor genes *ceh-34, tbx-2, ceh-20, eya-1, hlh-1* and the muscle structure genes *deb-1, unc-78, unc-112, sgn-1, tmd-2* throughout a 40 minute transcription inhibition time course in muscle cells. Each point represents normalized transcript abundance from one of three biological replicates. (F) Box plots showing the neuron-specific mRNA half-life distributions of neural fate-specifying transcription factor genes, synapse-/dendrite-/axon-associated genes, and all other genes. (G, H) Scatter plots of the normalized transcript abundance of the neural fate-specifying transcription factor genes *ceh-5, egl-46, unc-86, ceh-43, hlh-2* and the synapse-/dendrite-/axon-associated genes *lnp-1, unc-37, mig-2, ced-10, unc-53* throughout a 40 minute transcription inhibition time course in neuronal cells. Each point represents normalized transcript abundance from one of three biological replicates. Numbers to the left of the box plots are median half-lives within each group. Numbers above box plots are the number of genes with half-lives greater than 150 minutes within each group. P-value comparing median half-lives was calculated using the Wilcoxon rank sum test.

To determine what may drive these differences in half-life distributions, we examined the mRNA half-lives of genes that fell within specific functional categories in muscle and neuronal cells (**Supplemental Figure S11A-S11B**). Within muscle cells, transcripts encoding transcription factors with roles in mesoderm or muscle fate specification (such as *ceh-34, tbx-2, ceh-20, eya-1*, and *hlh-1*) had significantly shorter half-lives overall compared to transcripts encoding muscle structural proteins (such as *deb-1, unc-78, unc-112, sgn-1, tmd-2*) (**Figure 7C-7E**). Similarly, in neurons, transcripts encoding neural fate transcription factors (such as *ceh-5, egl-46, unc-86, ceh-43*, and *hlh-2*) had significantly shorter half-lives compared to transcripts associated with synapses, dendrites, or axons (such as *lnp-1, unc-37, mig-2, ced-10*, and *unc-53*) **(Figure 7F-7H**). While the overall mRNA half-life distributions for cell type-specific and broadly expressed genes were similar within epidermis and pharynx, in these cell classes we similarly saw faster decay of enriched transcription factor transcripts and slower decay of structural transcripts such as those encoding cuticle-associated proteins in the epidermis and peptidase inhibitors in the pharynx (**Supplemental Figure S10B-S10G, S11C-S11D**).

### Differential mRNA decay occurs across germline and somatic cells in the C. elegans embryo

The most striking cell type-specific difference in mRNA degradation was between the germline (median t_1/2_ = 62 minutes) and soma (median t_1/2_ = 36 minutes) (**Figures 4E, 8A**). When comparing mRNA half-lives of shared genes between these cell classes, about 80% of transcripts were more stable in the germline than they were in the soma (**Figure 8B**).

**Figure 8.**
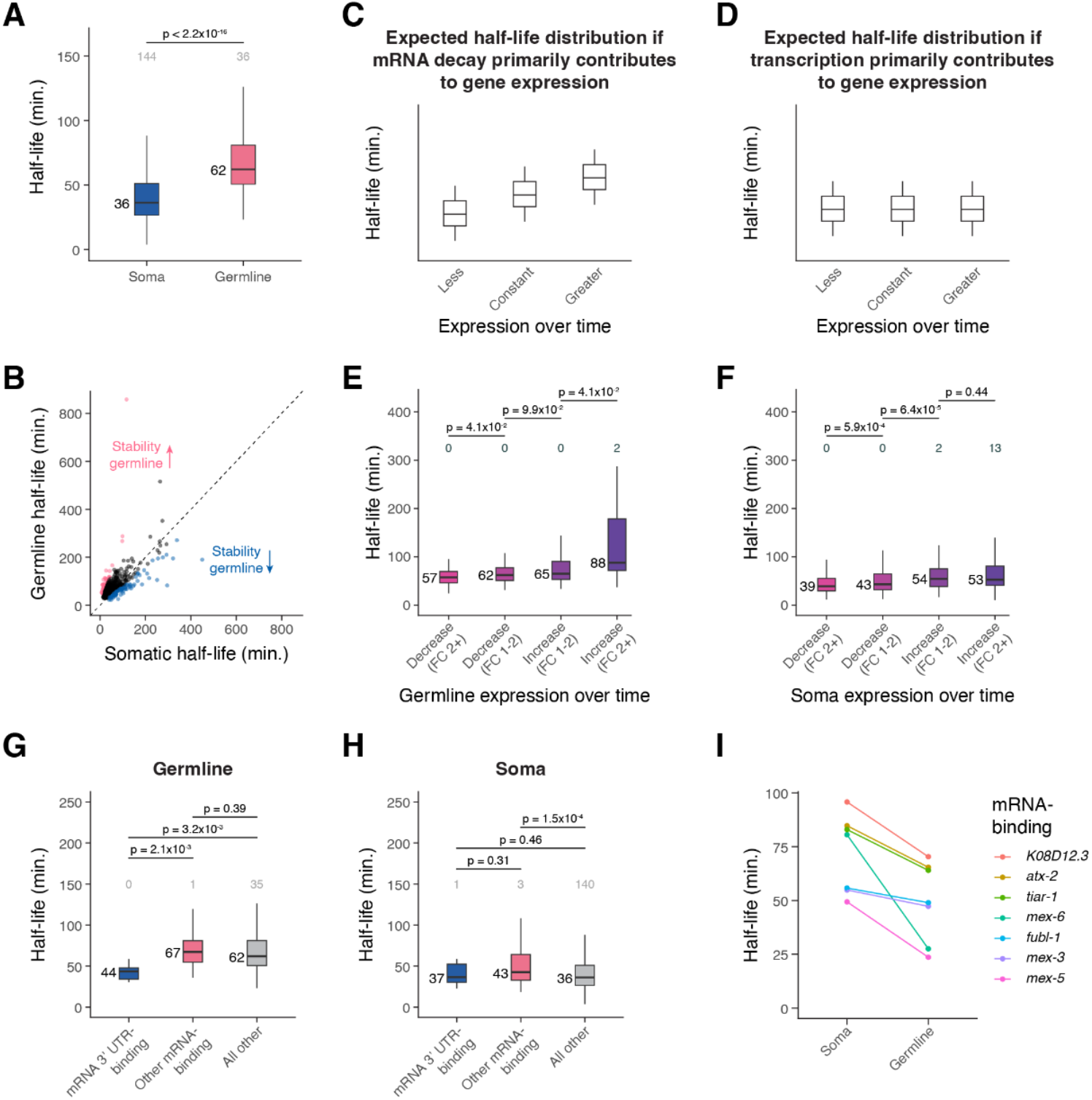
Differential mRNA decay occurs across germline and somatic cells in the *C. elegans* embryo. (A) Box plots showing the cell type-specific mRNA half-life distributions of genes across the germline and soma. (B) Scatter plot comparing soma- and germline-specific mRNA half-lives to one another. Each point represents a gene. Pink points correspond to the top 10% of genes with longer mRNA decay in the germline compared to in the soma. Blue points correspond to the top 10% of genes with faster mRNA decay in the germline compared to in the soma. Dashed line is the x = y line. (C) A cartoon of box plots representing the expected mRNA half-life distribution of genes with less, constant, or greater expression over time if mRNA decay primarily contributes to gene expression. (D) A cartoon of box plots representing the expected mRNA half-life distribution of genes with less, constant, or greater expression over time if transcription primarily contributes to gene expression. (E) Box plots showing the germline-specific mRNA half-life distributions for genes that decrease or increase in expression over time in the germline from a fold-change of 1-2 or greater than 2. (F) Box plots showing the soma-specific mRNA half-life distributions for genes that decrease or increase in expression over time in the soma from a fold-change of 1-2 or greater than 2. (G) Box plots showing the germline-specific mRNA half-life distributions of mRNA 3’ UTR-binding genes, other mRNA-binding genes, and genes not annotated as RNA-binding. (H) Box plots showing the soma-specific mRNA half-life distributions of mRNA 3’ UTR-binding genes, other mRNA-binding genes, and genes not annotated as RNA-binding. (I) Plot displaying the soma- and germline-line specific mRNA half-lives for mRNA-binding genes whose decay is more rapid in the germline than in the soma. Numbers to the left of the box plots are median half-lives within each group. Numbers above the box plots are the number of genes with half-lives greater than 150 minutes (A, G, H) or 325 minutes (E, F) within each group. P-value comparing median half-lives was calculated using the Wilcoxon rank sum test.

In the *C. elegans* embryo, the germline is derived from the P4 blastomere, which is born at the 24-cell stage after a series of asymmetric cell divisions (Wang and Seydoux 2013). At the 88-cell stage, P4 divides into the primordial germ cells Z2 and Z3, which do not divide further throughout embryogenesis. P4, Z2 and Z3 are thought to be transcriptionally quiescent (Wang and Seydoux 2013). Despite this, our *C. elegans* embryo single-cell atlas (Packer et al. 2019) identified early, mid-embryo, and late germline cells that had substantial quantitative differences in gene expression despite the supposed lack of transcription. This raises the question of whether differences in mRNA stability govern the quantitative maturation of the germline transcriptome during embryogenesis.

To test this, we compared mRNA stability between transcripts that increase or decrease in relative abundance across time in the germline. If expression differences result from differential mRNA decay, we would expect genes with decreasing relative abundance to have shorter mRNA half-lives compared to genes with constant expression (**Figure 8C**). Similarly, we would expect genes with increasing relative abundance to have longer mRNA half-lives. On the other hand, if changes in relative abundance result from previously unrecognized transcription, we might expect comparable half-life distributions for each group (**Figure 8D**).

We found that changes in germline expression over time correlated with mRNA stability, consistent with a role for mRNA turnover in sculpting the germline transcriptome. Transcripts that decrease in abundance over time had shorter half-lives overall, and transcripts that increase in abundance over time had longer half-lives overall (**Figure 8E**). This effect was stronger than the increase in mRNA half-lives we saw for genes with greater soma-specific expression over time (**Figure 8F**). These results are consistent with a model where changes in mRNA levels throughout embryogenesis result from mRNA stability differences in the germline and from a mix of transcriptional and post-transcriptional mechanisms in the soma.

The changes observed in transcript abundance in the germline raise the question of what types of genes increase or decrease over time. Gene ontology analysis revealed that germline-specific genes were enriched for genes associated with mRNA 3’ UTR-binding (**Supplemental Figure S11E**). We found that genes with this annotation had significantly shorter mRNA half-lives in the germline compared to those of other mRNA-binding genes or genes not annotated as RNA-binding (**Figure 8G**). This was not the case in the soma (**Figure 8H**), suggesting that germline maturation might be specifically regulated by decay of transcripts encoding RNA-binding proteins. Transcripts encoding mRNA-binding proteins with faster decay in the germline compared to in the soma included *mex-5* and *mex-6* transcripts, for which half-lives in the soma were more than twice as long as they were in the germline (**Figure 8I**). MEX-5 and MEX-6 are nearly identical proteins that are known to inhibit the translation of at least some germline proteins, and they become more highly expressed in somatic cells versus germline cells over time in the early embryo (Schubert et al. 2000). Our results suggest that maturation of the germline in part results from germline-specific turnover of maternally provided somatic regulators like *mex-5* and *mex-6*.

## Discussion

Using a transcription inhibition approach and RNA-sequencing, we measured mRNA half-lives transcriptome-wide throughout *C. elegans* embryogenesis. We found that global mRNA decay rates differ between cell types and developmental stages, identified gene attributes and sequence motifs associated with different rates of decay, and showed that regulated RNA turnover is associated with dynamic gene expression patterns. Our findings emphasize that developmental gene expression patterns result from regulation of both transcription and RNA decay.

A major advantage to the transcription inhibition approach is that it can be readily applied to any RNA-sequencing platform, and the downstream analyses to calculate mRNA half-lives transcriptome-wide are relatively straightforward in a dynamic situation like early embryogenesis. In addition, general concordance in measured mRNA decay rates has been observed between transcription inhibition approaches and approaches that maintain transcription, such as metabolic labeling, within mammalian cells and fly embryos (Burow et al. 2015; Herzog et al. 2017). Our analyses suggest that mRNA half-lives in the *C. elegans* embryo range from ∼20 to 200 minutes for most genes. Previous studies in cell culture systems found that median half-life is roughly proportional to cell cycle length (Vejnar et al. 2019). A study of decay in early *Drosophila melanogaster* embryos (nc11 through cephalic furrow formation), which have comparable cell cycle lengths (∼10-180 minutes) to those in the *C. elegans* embryo (Foe 1989; Bao et al. 2008), observed a similar range of half-lives to ours for ∼260 zygotic transcripts (Beadle et al. 2023). In older fly embryos (stage 12 to 15), which have longer cell cycles, the median half-life was longer at 73 minutes (Burow et al. 2015). This was longer than our *C. elegans* median half-life of 54 minutes, perhaps reflecting that older fly embryos no longer have the short mitotic cycles and rapid development characteristic of early fly embryos. We similarly found that mRNA half-lives increase over time during *C. elegans* embryogenesis. It is important to emphasize that some genes go against this trend, however, and have faster mRNA decay at later embryonic stages. The fact that these genes are enriched for specific categories of genes, such as transcription factor genes or genes involved in building the cilium, suggests that the rate of RNA turnover across time is heavily regulated.

While these changes are consistent with a relationship between the cell cycle and RNA turnover, they could also reflect the need for progenitor cells to degrade mRNAs in a timely manner to allow cell differentiation to proceed appropriately, as has been observed in other systems (Soustelle et al. 2008; Abbadi et al. 2019). Consistent with this, we observed that mRNAs encoding transcription factors had faster mRNA degradation compared to other transcripts, and fast RNA turnover was significantly associated with transient transcription factor protein expression. These results highlight that mRNA degradation likely plays a key role in modulating the protein expression of developmental regulators.

We found that specific functional categories of genes had distinct mRNA decay dynamics in different cell types. For example in neurons, transcripts encoding neural fate-specifying transcription factors had relatively short half-lives, while those associated with synapses, dendrites, or axons had relatively long half-lives. Similar findings were observed for neuron-specific mRNA half-lives in the fly embryo (Burow et al. 2015). Perhaps the unique architecture of neuronal cells can help explain these differences in the regulation of mRNA degradation. If transcripts are required distant from the cell body at axons and dendrites, their stabilization may facilitate localized translation. We also found that transcripts vary substantially in their germline mRNA stability, and this is correlated with changes in germline expression over time. This suggests that the regulation of mRNA degradation plays an important role in shaping the germline transcriptome. The relatively rapid decay of transcripts encoding mRNA 3’ UTR-binding proteins in the germline further points to the importance of post-transcriptional regulation in this cell type.

From yeast to human cells, greater translational efficiency is associated with greater transcript stability (Presnyak et al. 2015; Wu et al. 2019). Our observation that stable transcripts had significantly higher optimal codon usage compared to unstable transcripts in *C. elegans* embryos is consistent with this. We also found that stable transcripts tend to have more introns than unstable transcripts, in line with the presence of introns having a stabilizing effect in mammals and plants (Shaul 2017) and increasing protein production in *C. elegans* (Crane et al. 2019). The relationship between 3’ UTR length and mRNA stability appears to differ depending on the context. In zebrafish, longer 3’ UTRs conferred resistance to codon-mediated deadenylation for maternal transcripts (Mishima and Tomari 2016). Our data align with this finding, as stable transcripts had significantly longer 3’ UTRs compared to unstable transcripts. Future studies examining the effect of altering such features will test the potentially combinatorial causal relationships between these features and mRNA stability.

Examining the mRNA half-lives of genes that accumulate to high transcript levels revealed that such genes undergo overall slower transcript turnover compared to genes that accumulate to low transcript levels. Some genes accumulate so rapidly in early embryogenesis that their transcription rates must be high, irrespective of their mRNA turnover rates (Sivaramakrishnan et al. 2023). This emphasizes that the regulation of both transcription and mRNA degradation can work together to achieve high gene expression. The 3’ UTRs of genes with high transcript accumulation were significantly enriched for several motifs, including poly(C) motifs. In mammals, poly(C)-binding proteins have been implicated in the stabilization of transcripts (Makeyev and Liebhaber 2002), and transcripts that contain a poly(C) motif may be stabilized in *C. elegans* oocytes (Stoeckius et al. 2014). Future studies will establish whether poly(C)-binding protein homologs act as transcript stabilizers in the *C. elegans* embryo.

## Methods

### Embryonic cell isolation and culturing

*C. elegans* adults (N2 strain) were grown on large, enriched peptone plates seeded with NA22 bacteria. Mixed-stage embryos were released from adult worms using hypochlorite treatment followed by two washes with previously described complete L-15 cell culture media (Bianchi and Driscoll 2006). To obtain cell suspensions, embryos were treated with 0.5 mg/ml chitinase in egg buffer on ice until the eggshells were dissolved (about 10 minutes). Then embryonic cells were dissociated using a 3 ml syringe fitted with 21 1/2 gauge needle until >80% of embryos were disrupted. The cell suspension was passed through a 10 μM filter before being washed then resuspended in complete L-15.

### Bulk RNA-sequencing transcription inhibition time course

Following embryonic cell isolation, cells were treated with trypan blue and counted using a hemocytometer. Cell cultures at a concentration of ∼1 million cells/ml were treated with 2 μg/ml of the transcription inhibitor actinomycin D (actD) over a time course of 0, 10, 20, 40, and 60 minutes at 20 degrees C. Each time point had 2 ml of cell suspension (∼2 million cells). After each time point, RNA was isolated from cells using QIAzol that contained a 1:350,000 dilution of the ERCC spike-in kit (Baker et al. 2005). Libraries were prepared using the Illumina Ribo-Zero Plus rRNA Depletion Kit and sequenced on an Illumina NextSeq 500.

### Bulk RNA-sequencing computational analysis

The RNA-sequencing data were processed using the STAR (Spliced Transcripts Alignment to a Reference) alignment tool (Dobin et al. 2013) against the WormBase WS277 reference transcriptome with 3’ UTR extensions as previously carried out (Packer et al. 2019). VERSE was used to get read counts for genes (Zhu et al. 2016). Genes had to have a count greater than 30 at the 0 minute time point across all biological replicates for their expression data to be used in downstream analyses. For half-life comparisons between biological replicates, only well-measured genes were examined. These were genes with a count greater than 30 at the 0 minute time point whose expression throughout the transcription inhibition time course fit an exponential decay model R^2^ ≥ 0.75 within each replicate.

### Single-cell RNA-sequencing transcription inhibition time course

Cell cultures at a concentration of ∼1 million cells/ml were treated with 2 μg/ml of the transcription inhibitor actinomycin D (actD) over a time course of 0, 20, and 40 minutes at 20 degrees C. Each time point had 2 ml of cell suspension (∼2 million cells). After each time point, cells were kept on ice. At the end of the 40 minute time course, cells were washed with complete L-15, washed with 1X PBS with 0.4% BSA, then resuspended in 1X PBS with 0.4% BSA. Single cell capture and library preparation followed 10X Genomics published protocols for the Chromium Next GEM Single Cell 3’ Reagent Kits v3.1, with the objective of recovering ∼10,000 cells for each time point. Libraries were sequenced on an Illumina NextSeq 500.

### Single-cell RNA-sequencing computational analysis

The data were processed with the 10X Genomics CellRanger pipeline, aligning reads to the WormBase WS277 reference transcriptome with 3’ UTR extensions as previously carried out(Packer et al. 2019). Ambient RNA contamination was removed using the SoupX algorithm (Young and Behjati 2020) with default settings. The data were then visualized using dimensionality reduction methods for either merged datasets within each biological replicate or an integrated dataset of all biological replicates. Single-cell transcriptomes were first projected into 150 dimensions using PCA and then projected into two dimensions using the UMAP algorithm. For each cell, the age of the embryo from which it came from was estimated by correlating its transcriptome with a whole-embryo bulk RNA-seq time series as previously described (Hashimshony et al. 2015). Cells were then automatically annotated with their corresponding cell type or manually annotated based on cell type-specific marker genes reported in WormBase (Lee et al. 2018). Automated annotation was done using Seurat to project the PCA structure of a reference, an existing *C. elegans* embryo single cell atlas (Packer et al. 2019), to our single-cell data. Epidermal cells were manually annotated to better include progenitor cells that the reference dataset did not include. Within the cell types we examined, doublets were manually removed. This was done by identifying clusters of doublets in iterated UMAP projections of the data on the basis of co-expression of cell type-specific marker genes. Genes with a UMI count greater than 30 at the zero minute time point combined across all biological replicates were used in downstream analyses. For half-life comparisons between biological replicates, only well-measured genes were examined. These were genes with an UMI greater than 30 at the 0 minute time point whose expression throughout the transcription inhibition time course fit an exponential decay model R^2^ ≥ 0.75 within each replicate.

### mRNA half-life calculations

For the bulk RNA sequencing data, gene counts were normalized to the sum of counts corresponding to ERCC RNAs. Normalized gene counts were then fit to an exponential decay model (A(t) = A_0_e^-kt^) using nonlinear least squares regression in R, allowing a decay rate constant (k_decay_) to be calculated for each gene. A half-life was then calculated for each gene using the following equation: t_1/2_ = ln(2)/k_decay_. 95% confidence intervals for k_decay_ were determined using the R package nlstools. For the single-cell RNA sequencing data, half-lives were calculated in a similar manner with the number of UMIs per gene. Transcript abundance was normalized to the sum of UMIs corresponding to ribosomal protein genes, and this normalized expression was further adjusted by a correction factor to account for the decay of ribosomal protein genes over time (0.5^time point/median ribosomal protein gene half-life^, where median ribosomal protein gene half-life was determined from the bulk RNA-sequencing data).

A combination of coefficient of variation and confidence intervals was used to filter out poorly measured half-lives. Gene half-lives were included in downstream analyses if their coefficient of variation (standard deviation/mean^*^100) across biological replicates was ≤ 50% or the fold-change between the upper limit of its 95% confidence interval and measured half-life was ≤ 3. To better include high-stability mRNAs, genes with half-lives > 100 minutes were allowed a looser filtering strategy. Such genes were kept if their half-lives had a coefficient of variation ≤ 75% or fold-change between the upper limit of its 95% confidence interval and measured half-life ≤ 4.

### Gene ontology analysis

Gene ontology analysis was performed using the WormBase Enrichment Analysis tool (Angeles-Albores et al. 2018) against relevant background sets of genes. For example, for the bulk RNA sequencing data examining the top 15% slow- and fast-decaying genes, the background set of genes used was all genes that met our moderate half-life filtering metric.

### Transcript sequence feature analysis

3’ UTRs across the *C. elegans* transcriptome were retrieved from the WormBase ParaSite database (Howe et al. 2017) on March 9, 2023. Only 3’ UTRs with ≥ 8 nucleotides were used for downstream analyses. To examine percent GC content, the longest 3’ UTR was used for genes with multiple isoforms. To examine the correlation between 3’ UTR length and half-life, mean UTR length across isoforms for each gene was calculated.

Coding sequences across the *C. elegans* transcriptome were retrieved from the WormBase ParaSite database(Howe et al. 2017) on March 4, 2023. To examine percent GC content, the longest coding sequence was used for genes with multiple isoforms. To examine the correlation between coding sequence length and half-life, mean coding sequence length across isoforms for each gene was calculated. *C. elegans*-specific tRNA adaptation index (tAI) weights were used from stAIcalc (Sabi et al. 2017) (http://tau-tai.azurewebsites.net/) to calculate a tAI value for each gene, with the longest coding sequence being used for genes with multiple isoforms. The tAI value for each gene was the geometric mean of tAI weights for all codons in the coding sequence (Yoon et al. 2018).

Introns were identified using the TxDb.Celegans.UCSC.ce11.refGene annotation package for TxDb objects. To examine the correlation between the number of introns and half-life, the mean number of introns across isoforms for each gene was calculated.

The ‘regsubsets’ function from the R package ‘leaps’ was used for linear multiple regression analysis. mRNA half-life was the modeled variable and the sequence features of interest (tAI, mean intron, mean 3’ UTR length, mean coding sequence length, GC content in 3’ UTR, GC content in coding sequence) the predictors. The individual contribution of each sequence feature to variation in half-lives was evaluated by having each feature as the only predictor in the model.

### Motif analysis

MEME differential enrichment was used for motif analysis (https://meme-suite.org/meme/tools/meme). The minimum width for motifs was set to 5 and the maximum width for motifs was set to 15. For motif analysis in 3’ UTRs, the longest 3’ UTR was used for genes with multiple isoforms. To identify motifs in the 3’ UTRs of the top 15% slow-decaying genes in our bulk data, the 3’ UTRs of the top 15% fast-decaying genes were used as control sequences (and vice versa).

### Determining transient or persistent gene expression

We identified genes with highly transient, transient, persistent or highly persistent expression over time using our previously published *C. elegans* embryo single cell atlas (Packer et al. 2019). Highly transient genes were defined as genes whose expression was lost from parent to daughter cell in ≥ 30 comparisons or ≥ 30% of comparisons. Transient genes were defined as genes whose expression was lost from parent to daughter cell in ≥ 10 comparisons or ≥ 10% of comparisons, excluding highly transient genes. Highly persistent genes were defined as genes whose expression was maintained from parent to daughter cell in ≥ 30 comparisons or ≥ 30% of comparisons. Furthermore, these genes had to have ≤ 1 comparison and ≤ 1% of comparisons with a loss in expression from parent to daughter. Persistent genes were defined as genes whose expression was maintained from parent to daughter cell in ≥ 10 comparisons or ≥ 10% of comparisons, excluding highly persistent genes. Furthermore, these genes had to have ≤ 5 comparisons and ≤ 5% of comparisons with a loss in expression from parent to daughter.

### Determining transient or persistent protein expression for transcription factors

To characterize transcription factor protein expression dynamics, we used expression data provided from a single-cell transcription factor protein expression atlas of the *C. elegans* embryo (Ma et al. 2021). A transcription factor was considered expressed in a cell if its expression was > 1.5^*^IQR below the first quartile for protein expression across all cells. Transient proteins were defined as proteins whose expression was lost from parent to daughter cell in ≥ 10 comparisons or ≥ 10% of comparisons. Persistent proteins were defined as proteins whose expression was maintained from parent to daughter cell in ≥ 10 comparisons or ≥ 10% of comparisons. Proteins with data from more than one reporter strain were only characterized as transient or persistent if the strains agreed with one another.

### Hierarchical clustering of gene expression patterns

Gene expression was first scaled using the ‘scale’ function in R, then a distance metric was calculated between each pair of genes using the ‘dist’ function. Hierarchical clustering was performed on these distance values using the ‘hclust’ function. Cluster number was chosen based on visualization with a dendrogram.

### Determining genes with high transcript accumulation

To characterize genes in the staged embryo RNA-sequencing data (Hashimshony et al. 2015) whose expression peaks around 200 minutes past the four-cell embryo stage as having high, medium, or low transcript accumulation, we calculated a slope value for each gene. This slope = (maximum expression from embryo stages 10 to 200 minutes - minimum expression from embryo stages 10 to 200 minutes) / maximum stage in minutes - minimum stage in minutes. Slopes from 0 to 0.2 quantile were considered low-accumulating genes, slopes from 0.4 to 0.6 quantile were considered medium-accumulating genes, and slopes from 0.8 to 1.0 quantile were considered high-accumulating genes. A similar analysis was done for genes whose expression peaks around 350 minutes past the four-cell embryo stage.

### Determining cell type-specific and broadly expressed genes

Data from our *C. elegans* embryo single cell atlas (Packer et al. 2019) was used to determine gene expression within muscle, germline, epidermis, neuron, and pharynx cells in transcripts per million (TPM). A gene was considered to be specific to a given cell type if it was expressed more than twofold greater in that cell type compared to all other cell types. For example, a gene was considered muscle-specific if it was expressed more than twofold greater in muscle than it was in germline, epidermis, neuron, and pharynx. All other genes were considered broadly expressed. 10 TPM was added to each gene to avoid issues in comparing unexpressed or poorly measured genes.

### Determining zygotic-only genes

Zygotic-only genes were determined using a single-cell RNA-sequencing dataset of the early *C. elegans* embryo (Tintori et al. 2016). Genes with an average reads per kilobase million (RPKM) > 50 within one-cell embryos were considered to be maternally-expressed genes. All other genes were considered to be zygotic-only genes.

## Data access

All RNA-sequencing data generated in this study have been submitted to the NCBI BioProject database (https://www.ncbi.nlm.nih.gov/bioproject) under accession number PRJNA1061171. R codes regarding mRNA half-life calculations for both the bulk and single-cell RNA-sequencing datasets are available at GitHub (https://github.com/fe-peng/celegans_embryo_mRNA_decay).

## Acknowledgements

We thank members of the Murray lab, the Penn Worm Group, Naveen Jain (University of Pennsylvania, Arjun Raj lab), Calvin Huang (University of California, Davis, Bo Liu lab), and Chenxin Li (University of Georgia, Robin Buell lab) for providing valuable discussion and comments on the manuscript. We also thank Jean Rosario and Catherine Lucey (University of Pennsylvania, Junhyong Kim lab) for their help in sequencing RNA libraries. This work was funded by R35GM127093, F31HD10785, and T32GM008216.

## Author contributions

F.P. and J.I.M. conceived and designed the study; F.P. performed the experiments; C.E.N. created new genome and transcriptome references and processed the bulk and single-cell RNA-sequencing data; F.P. performed analyses and visualization; J.I.M. supervised the analyses; F.P. wrote the original draft; F.P., C.E.N., and J.I.M. reviewed and edited the manuscript.

## References

Abbadi D, Yang M, Chenette DM, Andrews JJ, Schneider RJ. 2019. Muscle development and regeneration controlled by AUF1-mediated stage-specific degradation of fate-determining checkpoint mRNAs. Proc Natl Acad Sci. 116(23):11285–11290. doi:10.1073/pnas.1901165116.

Alonso CR. 2012. A complex ‘mRNA degradation code’ controls gene expression during animal development. Trends Genet. 28(2):78–88. doi:10.1016/j.tig.2011.10.005.

Angeles-Albores D, Lee RYN, Chan J, Sternberg PW. 2018 Mar 2. Two new functions in the WormBase Enrichment Suite. MicroPublication Biol. doi:10.17912/W25Q2N. [accessed 2023 Apr 13]. https://www.micropublication.org/journals/biology/w25q2n.

Bae H, Coller J. 2022. Codon optimality-mediated mRNA degradation: Linking translational elongation to mRNA stability. Mol Cell. 82(8):1467–1476. doi:10.1016/j.molcel.2022.03.032.

Bailey TL, Johnson J, Grant CE, Noble WS. 2015. The MEME Suite. Nucleic Acids Res. 43(Web Server issue):W39–W49. doi:10.1093/nar/gkv416.

Baker SC, Bauer SR, Beyer RP, Brenton JD, Bromley B, Burrill J, Causton H, Conley MP, Elespuru R, Fero M, et al. 2005. The External RNA Controls Consortium: a progress report. Nat Methods. 2(10):731–734. doi:10.1038/nmeth1005-731.

Bao Z, Zhao Z, Boyle TJ, Murray JI, Waterston RH. 2008. Control of Cell Cycle Timing during C. elegans Embryogenesis. Dev Biol. 318(1):65–72. doi:10.1016/j.ydbio.2008.02.054.

Beadle LF, Love JC, Shapovalova Y, Artemev A, Rattray M, Ashe HL. 2023. Combined modelling of mRNA decay dynamics and single-molecule imaging in the Drosophila embryo uncovers a role for P-bodies in 5′ to 3′ degradation. PLOS Biol. 21(1):e3001956. doi:10.1371/journal.pbio.3001956.

Becht E, McInnes L, Healy J, Dutertre C-A, Kwok IWH, Ng LG, Ginhoux F, Newell EW. 2019. Dimensionality reduction for visualizing single-cell data using UMAP. Nat Biotechnol. 37(1):38–44. doi:10.1038/nbt.4314.

Bhattacharyya SN, Habermacher R, Martine U, Closs EI, Filipowicz W. 2006. Relief of microRNA-mediated translational repression in human cells subjected to stress. Cell. 125(6):1111–1124. doi:10.1016/j.cell.2006.04.031.

Bianchi L, Driscoll M. 2006. Culture of embryonic C. elegans cells for electrophysiological and pharmacological analyses. WormBook. [accessed 2023 Apr 12]. https://www.ncbi.nlm.nih.gov/books/NBK19713/.

Blacque OE, Li C, Inglis PN, Esmail MA, Ou G, Mah AK, Baillie DL, Scholey JM, Leroux MR. 2006. The WD Repeat-containing Protein IFTA-1 Is Required for Retrograde Intraflagellar Transport. Mol Biol Cell. 17(12):5053–5062. doi:10.1091/mbc.E06-06-0571.

Brocal-Ruiz R, Esteve-Serrano A, Mora-Martínez C, Franco-Rivadeneira ML, Swoboda P, Tena JJ, Vilar M, Flames N. 2023. Forkhead transcription factor FKH-8 cooperates with RFX in the direct regulation of sensory cilia in Caenorhabditis elegans. Portman D, Sengupta P, editors. eLife. 12:e89702. doi:10.7554/eLife.89702.

Burghoorn J, Dekkers MPJ, Rademakers S, de Jong T, Willemsen R, Jansen G. 2007. Mutation of the MAP kinase DYF-5 affects docking and undocking of kinesin-2 motors and reduces their speed in the cilia of Caenorhabditis elegans. Proc Natl Acad Sci U S A. 104(17):7157–7162. doi:10.1073/pnas.0606974104.

Burow DA, Umeh-Garcia MC, True MB, Bakhaj CD, Ardell DH, Cleary MD. 2015. Dynamic regulation of mRNA decay during neural development. Neural Develop. 10(1):11. doi:10.1186/s13064-015-0038-6.

Christensen M, Estevez A, Yin X, Fox R, Morrison R, McDonnell M, Gleason C, Miller DM, Strange K. 2002. A primary culture system for functional analysis of C. elegans neurons and muscle cells. Neuron. 33(4):503–514. doi:10.1016/s0896-6273(02)00591-3.

Crane MM, Sands B, Battaglia C, Johnson B, Yun S, Kaeberlein M, Brent R, Mendenhall A. 2019. In vivo measurements reveal a single 5′-intron is sufficient to increase protein expression level in Caenorhabditis elegans. Sci Rep. 9(1):9192. doi:10.1038/s41598-019-45517-0.

De-Castro ARG, Quintas-Gonçalves J, Silva-Ribeiro T, Rodrigues DRM, De-Castro MJG, Abreu CM, Dantas TJ. The IFT20 homolog in Caenorhabditis elegans is required for ciliogenesis and cilia-mediated behavior. MicroPublication Biol. 2021: 10.17912/micropub.biology.000396. doi:10.17912/micropub.biology.000396.

Dobin A, Davis CA, Schlesinger F, Drenkow J, Zaleski C, Jha S, Batut P, Chaisson M, Gingeras TR. 2013. STAR: ultrafast universal RNA-seq aligner. Bioinformatics. 29(1):15–21. doi:10.1093/bioinformatics/bts635.

Edgar LG. 1995. Blastomere culture and analysis. Methods Cell Biol. 48:303–321. doi:10.1016/s0091-679x(08)61393-x.

Efimenko E, Blacque OE, Ou G, Haycraft CJ, Yoder BK, Scholey JM, Leroux MR, Swoboda P. 2006. Caenorhabditis elegans DYF-2, an Orthologue of Human WDR19, Is a Component of the Intraflagellar Transport Machinery in Sensory Cilia. Mol Biol Cell. 17(11):4801–4811. doi:10.1091/mbc.E06-04-0260.

Foe VE. 1989. Mitotic domains reveal early commitment of cells in Drosophila embryos. Development. 107(1):1–22. doi:10.1242/dev.107.1.1.

Fuxman Bass JI, Pons C, Kozlowski L, Reece-Hoyes JS, Shrestha S, Holdorf AD, Mori A, Myers CL, Walhout AJ. 2016. A gene-centered C. elegans protein–DNA interaction network provides a framework for functional predictions. Mol Syst Biol. 12(10):884. doi:10.15252/msb.20167131.

Geuens T, Bouhy D, Timmerman V. 2016. The hnRNP family: insights into their role in health and disease. Hum Genet. 135(8):851–867. doi:10.1007/s00439-016-1683-5.

van der Giessen K, Gallouzi I-E. 2007. Involvement of Transportin 2–mediated HuR Import in Muscle Cell Differentiation. Mol Biol Cell. 18(7):2619–2629. doi:10.1091/mbc.E07-02-0167.

Giraldez AJ, Mishima Y, Rihel J, Grocock RJ, Van Dongen S, Inoue K, Enright AJ, Schier AF. 2006. Zebrafish MiR-430 promotes deadenylation and clearance of maternal mRNAs. Science. 312(5770):75–79. doi:10.1126/science.1122689.

Hashimshony T, Feder M, Levin M, Hall BK, Yanai I. 2015. Spatiotemporal transcriptomics reveals the evolutionary history of the endoderm germ layer. Nature. 519(7542):219–222. doi:10.1038/nature13996.

Haycraft CJ, Schafer JC, Zhang Q, Taulman PD, Yoder BK. 2003. Identification of CHE-13, a novel intraflagellar transport protein required for cilia formation. Exp Cell Res. 284(2):249–261. doi:10.1016/S0014-4827(02)00089-7.

Herzog VA, Reichholf B, Neumann T, Rescheneder P, Bhat P, Burkard TR, Wlotzka W, von Haeseler A, Zuber J, Ameres SL. 2017. Thiol-linked alkylation of RNA to assess expression dynamics. Nat Methods. 14(12):1198–1204. doi:10.1038/nmeth.4435.

Howe KL, Bolt BJ, Shafie M, Kersey P, Berriman M. 2017. WormBase ParaSite − a comprehensive resource for helminth genomics. Mol Biochem Parasitol. 215:2–10. doi:10.1016/j.molbiopara.2016.11.005.

Jiang P, Singh M, Coller HA. 2013. Computational assessment of the cooperativity between RNA binding proteins and MicroRNAs in Transcript Decay. PLoS Comput Biol. 9(5):e1003075. doi:10.1371/journal.pcbi.1003075.

Kobayashi T, Gengyo-Ando K, Ishihara T, Katsura I, Mitani S. 2007. IFT-81 and IFT-74 are required for intraflagellar transport in C. elegans. Genes Cells Devoted Mol Cell Mech.12(5):593–602. doi:10.1111/j.1365-2443.2007.01076.x.

Kunitomo H, Iino Y. 2008. Caenorhabditis elegans DYF-11, an orthologue of mammalian Traf3ip1/MIP-T3, is required for sensory cilia formation. Genes Cells. 13(1):13–25. doi:10.1111/j.1365-2443.2007.01147.x.

Łabno A, Tomecki R, Dziembowski A. 2016. Cytoplasmic RNA decay pathways - Enzymes and mechanisms. Biochim Biophys Acta BBA - Mol Cell Res. 1863(12):3125–3147. doi:10.1016/j.bbamcr.2016.09.023.

Lambacher NJ, Bruel A-L, van Dam TJP, Szymańska K, Slaats GG, Kuhns S, McManus GJ, Kennedy JE, Gaff K, Wu KM, et al. 2016. TMEM107 recruits ciliopathy proteins to subdomains of the ciliary transition zone and causes Joubert syndrome. Nat Cell Biol. 18(1):122–131. doi:10.1038/ncb3273.

Lee RYN, Howe KL, Harris TW, Arnaboldi V, Cain S, Chan J, Chen WJ, Davis P, Gao S, Grove C, et al. 2018. WormBase 2017: molting into a new stage. Nucleic Acids Res. 46(Database issue):D869–D874. doi:10.1093/nar/gkx998.

Li C, Jensen VL, Park K, Kennedy J, Garcia-Gonzalo FR, Romani M, Mori RD, Bruel A-L, Gaillard D, Doray B, et al. 2016. MKS5 and CEP290 Dependent Assembly Pathway of the Ciliary Transition Zone. PLOS Biol. 14(3):e1002416. doi:10.1371/journal.pbio.1002416.

Li W, Yi P, Ou G. 2015. Somatic CRISPR–Cas9-induced mutations reveal roles of embryonically essential dynein chains in Caenorhabditis elegans cilia. J Cell Biol. 208(6):683–692. doi:10.1083/jcb.201411041.

Ma X, Zhao Z, Xiao L, Xu W, Kou Y, Zhang Y, Wu G, Wang Y, Du Z. 2021. A 4D single-cell protein atlas of transcription factors delineates spatiotemporal patterning during embryogenesis. Nat Methods. 18(8):893–902. doi:10.1038/s41592-021-01216-1.

Makeyev AV, Liebhaber SA. 2002. The poly(C)-binding proteins: a multiplicity of functions and a search for mechanisms. RNA. 8(3):265–278.

Mayr C. 2019. What Are 3′ UTRs Doing? Cold Spring Harb Perspect Biol. 11(10):a034728. doi:10.1101/cshperspect.a034728.

McInnes L, Healy J, Melville J. 2020. UMAP: Uniform Manifold Approximation and Projection for Dimension Reduction. doi:10.48550/arXiv.1802.03426. [accessed 2023 Oct 3]. http://arxiv.org/abs/1802.03426.

Mijalkovic J, Van Krugten J, Oswald F, Acar S, Peterman EJG. 2018. Single-Molecule Turnarounds of Intraflagellar Transport at the C. elegans Ciliary Tip. Cell Rep. 25(7):1701–1707.e2. doi:10.1016/j.celrep.2018.10.050.

Mishima Y, Tomari Y. 2016. Codon Usage and 3′ UTR Length Determine Maternal mRNA Stability in Zebrafish. Mol Cell. 61(6):874–885. doi:10.1016/j.molcel.2016.02.027.

Narsai R, Howell KA, Millar AH, O’Toole N, Small I, Whelan J. 2007. Genome-Wide Analysis of mRNA Decay Rates and Their Determinants in Arabidopsis thaliana. Plant Cell. 19(11):3418–3436. doi:10.1105/tpc.107.055046.

Nechipurenko IV, Sengupta P. 2017. The rise and fall of basal bodies in the nematode Caenorhabditis elegans. Cilia. 6(1):9. doi:10.1186/s13630-017-0053-9.

Ou G, Koga M, Blacque OE, Murayama T, Ohshima Y, Schafer JC, Li C, Yoder BK, Leroux MR, Scholey JM. 2007. Sensory Ciliogenesis in Caenorhabditis elegans: Assignment of IFT Components into Distinct Modules Based on Transport and Phenotypic Profiles. Mol Biol Cell. 18(5):1554–1569. doi:10.1091/mbc.E06-09-0805.

Packer JS, Zhu Q, Huynh C, Sivaramakrishnan P, Preston E, Dueck H, Stefanik D, Tan K, Trapnell C, Kim J, et al. 2019. A lineage-resolved molecular atlas of C. elegans embryogenesis at single cell resolution. Genomics. [accessed 2021 Mar 10]. http://biorxiv.org/lookup/doi/10.1101/565549.

Passmore LA, Coller J. 2021 Sep 30. Roles of mRNA poly(A) tails in regulation of eukaryotic gene expression. Nat Rev Mol Cell Biol.:1–14. doi:10.1038/s41580-021-00417-y.

Presnyak V, Alhusaini N, Chen Y-H, Martin S, Morris N, Kline N, Olson S, Weinberg D, Baker KE, Graveley BR, et al. 2015. Codon Optimality Is a Major Determinant of mRNA Stability. Cell. 160(6):1111–1124. doi:10.1016/j.cell.2015.02.029.

Qin H, Rosenbaum JL, Barr MM. 2001. An autosomal recessive polycystic kidney disease gene homolog is involved in intraflagellar transport in C. elegans ciliated sensory neurons. Curr Biol. 11(6):457–461. doi:10.1016/S0960-9822(01)00122-1.

Ray D, Kazan H, Cook KB, Weirauch MT, Najafabadi HS, Li X, Gueroussov S, Albu M, Zheng H, Yang A, et al. 2013. A compendium of RNA-binding motifs for decoding gene regulation. Nature. 499(7457):172–177. doi:10.1038/nature12311.

Roberson EC, Dowdle WE, Ozanturk A, Garcia-Gonzalo FR, Li C, Halbritter J, Elkhartoufi N, Porath JD, Cope H, Ashley-Koch A, et al. 2015. TMEM231, mutated in orofaciodigital and Meckel syndromes, organizes the ciliary transition zone. J Cell Biol. 209(1):129–142. doi:10.1083/jcb.201411087.

Sabi R, Volvovitch Daniel R, Tuller T. 2017. stAIcalc: tRNA adaptation index calculator based on species-specific weights. Bioinformatics. 33(4):589–591. doi:10.1093/bioinformatics/btw647.

Schubert CM, Lin R, Vries CJ de Plasterk RHA, Priess JR. 2000. MEX-5 and MEX-6 Function to Establish Soma/Germline Asymmetry in Early C. elegans Embryos. Mol Cell. 5(4):671–682. doi:10.1016/S1097-2765(00)80246-4.

Shaul O. 2017. How introns enhance gene expression. Int J Biochem Cell Biol. 91:145–155. doi:10.1016/j.biocel.2017.06.016.

Sivaramakrishnan P, Watkins C, Murray JI. 2023. Transcript accumulation rates in the early Caenorhabditis elegans embryo. Sci Adv. 9(34):eadi1270. doi:10.1126/sciadv.adi1270.

Soustelle L, Roy N, Ragone G, Giangrande A. 2008. Control of gcm RNA stability is necessary for proper glial cell fate acquisition. Mol Cell Neurosci. 37(4):657–662. doi:10.1016/j.mcn.2007.11.007.

Stoeckius M, Grün D, Kirchner M, Ayoub S, Torti F, Piano F, Herzog M, Selbach M, Rajewsky N. 2014. Global characterization of the oocyte-to-embryo transition in Caenorhabditis elegans uncovers a novel mRNA clearance mechanism. EMBO J. 33(16):1751–1766. doi:10.15252/embj.201488769.

Tadros W, Goldman AL, Babak T, Menzies F, Vardy L, Orr-Weaver T, Hughes TR, Westwood JT, Smibert CA, Lipshitz HD. 2007. SMAUG Is a Major Regulator of Maternal mRNA Destabilization in Drosophila and Its Translation Is Activated by the PAN GU Kinase. Dev Cell. 12(1):143–155. doi:10.1016/j.devcel.2006.10.005.

Thomsen S, Anders S, Janga SC, Huber W, Alonso CR. 2010. Genome-wide analysis of mRNA decay patterns during early Drosophiladevelopment. Genome Biol. 11(9):R93. doi:10.1186/gb-2010-11-9-r93.

Tintori SC, Osborne Nishimura E, Golden P, Lieb JD, Goldstein B. 2016. A Transcriptional Lineage of the Early C. elegans Embryo. Dev Cell. 38(4):430–444. doi:10.1016/j.devcel.2016.07.025.

Vastenhouw NL, Cao WX, Lipshitz HD. 2019. The maternal-to-zygotic transition revisited. Development. 146(11):dev161471. doi:10.1242/dev.161471.

Vejnar CE, Abdel Messih M, Takacs CM, Yartseva V, Oikonomou P, Christiano R, Stoeckius M, Lau S, Lee MT, Beaudoin J-D, et al. 2019. Genome wide analysis of 3’ UTR sequence elements and proteins regulating mRNA stability during maternal-to-zygotic transition in zebrafish. Genome Res. 29(7):1100–1114. doi:10.1101/gr.245159.118.

Wang JT, Seydoux G. 2013. Germ Cell Specification. Adv Exp Med Biol. 757:17–39. doi:10.1007/978-1-4614-4015-4_2.

Wang W, Jack BM, Wang HH, Kavanaugh MA, Maser RL, Tran PV. 2021. Intraflagellar Transport Proteins as Regulators of Primary Cilia Length. Front Cell Dev Biol. 9. [accessed 2023 Oct 10]. https://www.frontiersin.org/articles/10.3389/fcell.2021.661350.

Wu Q, Medina SG, Kushawah G, DeVore ML, Castellano LA, Hand JM, Wright M, Bazzini AA. 2019. Translation affects mRNA stability in a codon-dependent manner in human cells. Sonenberg N, Struhl K, Weissman JS, editors. eLife. 8:e45396. doi:10.7554/eLife.45396.

Yang E, van Nimwegen E, Zavolan M, Rajewsky N, Schroeder M, Magnasco M, Darnell JE. 2003. Decay Rates of Human mRNAs: Correlation With Functional Characteristics and Sequence Attributes. Genome Res. 13(8):1863–1872. doi:10.1101/gr.1272403.

Yoon J, Chung Y-J, Lee M. 2018. STADIUM: Species-Specific tRNA Adaptive Index Compendium. Genomics Inform. 16(4):e28. doi:10.5808/GI.2018.16.4.e28.

Young LE, Moore AE, Sokol L, Meisner-Kober N, Dixon DA. 2012. The mRNA stability factor HuR inhibits microRNA-16 targeting of COX-2. Mol Cancer Res MCR. 10(1):167–180. doi:10.1158/1541-7786.MCR-11-0337.

Young MD, Behjati S. 2020. SoupX removes ambient RNA contamination from droplet-based single-cell RNA sequencing data. GigaScience. 9(12):giaa151. doi:10.1093/gigascience/giaa151.

Zhu Q, Fisher SA, Shallcross J, Kim J. 2016. VERSE: a versatile and efficient RNA-Seq read counting tool. :053306. doi:10.1101/053306. [accessed 2023 Oct 6]. https://www.biorxiv.org/content/10.1101/053306v1.

