## Supplementary Figures for "A spatiotemporally resolved atlas of mRNA decay in the *C. elegans* embryo reveals differential regulation of mRNA stability across stages and cell types"

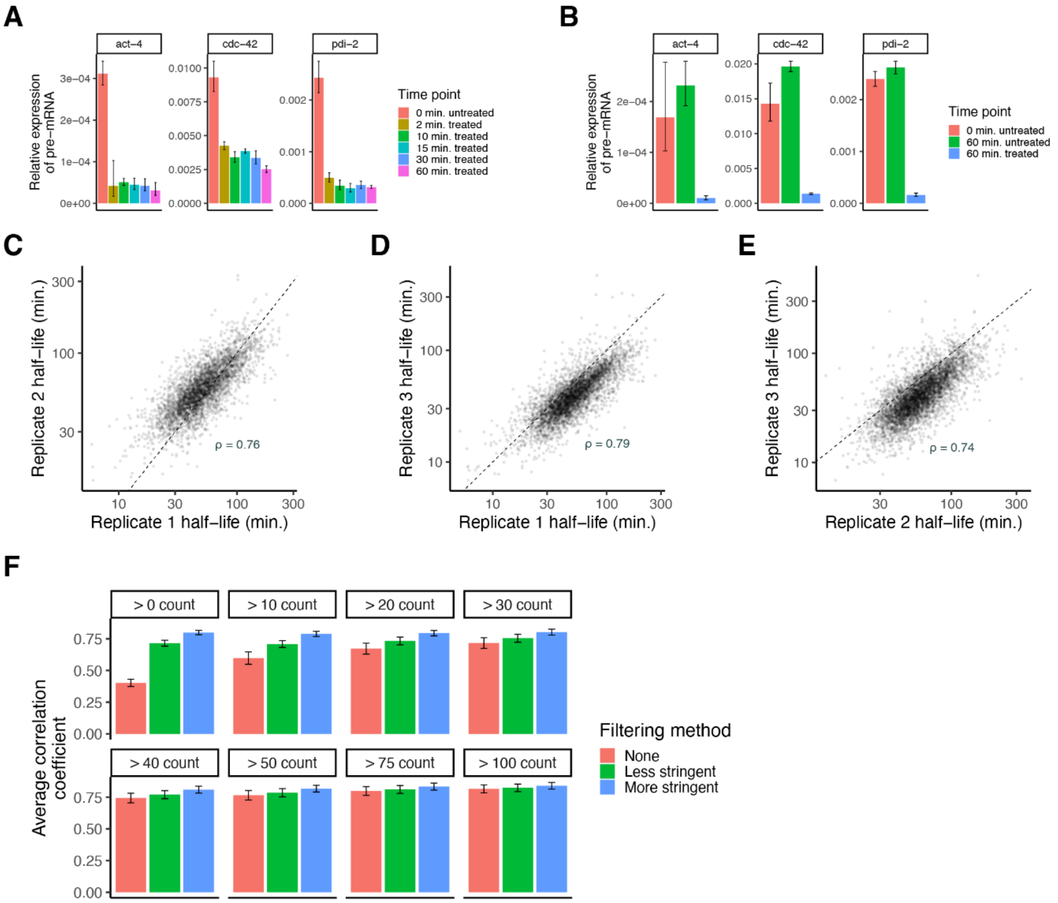


**Supplemental Figure S1. Quality control for transcription inhibition approach paired with bulk RNA-sequencing.** (A) Bar plots showing the relative expression of pre-mRNA for the housekeeping genes *act-4*, *cdc-42*, and *pdi-2* in embryonic cells following different lengths of transcription inhibition with actD. Expression was measured using RT-qPCR (quantitative reverse transcription PCR). Error bars represent variation in expression across three technical replicates. (B) Bar plots showing the relative expression of pre-mRNA for the housekeeping genes *act-4*, *cdc-42*, and *pdi-2* in embryonic cells following no treatment of actD, 60 minutes of treatment with actD, and 60 minutes with no treatment of actD. Expression was measured using RT-qPCR. Error bars represent variation in expression across three technical replicates. (C, D, E) Scatter plots showing the comparison of measured mRNA half-lives between three biological replicates on a log-log scale. Genes were compared if they had count > 30 at the 0 minute time point and if their decay fit an exponential decay model $R^2 \geq 0.75$ for each replicate. Spearman correlation coefficient is displayed for each pairwise comparison. Dashed line is the x = y line. (F) Bar plot showing the average Pearson correlation coefficient between mRNA half-lives across pairwise comparisons of three biological replicates under different count thresholds at the 0 minute time point and filtering methods. The less stringent filtering method only included genes if their coefficient of variation (standard deviation/mean*100) across biological replicates was ≤ 50% or the fold-change between the upper limit of their 95% confidence interval and measured half-life was ≤ 3. The stringent filtering method only included genes if their coefficient of variation across biological replicates was ≤ 30% or the fold-change between the upper limit of their 95% confidence interval and measured half-life was ≤ 2. Error bars represent standard deviation of the Pearson correlation coefficient among the three biological replicates.

**A**

| Term | Expected | Observed | Enrichment.Fold.Change | P.value | Q.value |
|---|---|---|---|---|---|
| cytosolic large ribosomal subunit GO:0022625 | 3.8 | 25 | 6.5 | 0.0e+00 | 0.0e+00 |
| extracellular region GO:0005576 | 21.0 | 62 | 2.9 | 7.2e−17 | 1.0e−14 |
| structural constituent of ribosome GO:0003735 | 14.0 | 47 | 3.4 | 1.3e−16 | 1.2e−14 |
| proton transmembrane transport GO:1902600 | 7.5 | 30 | 4.0 | 3.6e−14 | 2.5e−12 |
| supramolecular polymer GO:0099081 | 37.0 | 79 | 2.1 | 6.6e−12 | 3.7e−10 |
| calcium ion binding GO:0005509 | 15.0 | 41 | 2.8 | 3.0e−11 | 1.4e−09 |
| muscle system process GO:0003012 | 7.2 | 26 | 3.6 | 4.1e−11 | 1.7e−09 |
| structural constituent of chromatin GO:0030527 | 5.1 | 21 | 4.1 | 4.1e−11 | 1.7e−09 |
| transmembrane transport GO:0055085 | 58.0 | 105 | 1.8 | 1.2e−10 | 3.6e−09 |
| structural constituent of cytoskeleton GO:0005200 | 3.4 | 16 | 4.7 | 1.7e−10 | 4.9e−09 |
| myofibril GO:0030016 | 13.0 | 37 | 2.8 | 1.8e−10 | 4.9e−09 |
| actin binding GO:0003779 | 15.0 | 40 | 2.6 | 5.8e−10 | 1.4e−08 |
| transporter activity GO:0005215 | 57.0 | 102 | 1.8 | 6.3e−10 | 1.4e−08 |
| external encapsulating structure GO:0030312 | 7.1 | 24 | 3.4 | 1.1e−09 | 2.2e−08 |
| purine nucleoside triphosphate metabolic process GO:0009144 | 4.5 | 17 | 3.8 | 1.4e−08 | 2.6e−07 |
| protein heterodimerization activity GO:0046982 | 6.8 | 21 | 3.1 | 7.9e−08 | 1.4e−06 |
| peptide biosynthetic process GO:0043043 | 39.0 | 71 | 1.8 | 1.5e−07 | 2.5e−06 |
| muscle cell development GO:0055001 | 6.3 | 19 | 3.0 | 5.1e−07 | 8.1e−06 |
| actin filament−based process GO:0030029 | 26.0 | 50 | 1.9 | 1.0e−06 | 1.5e−05 |
| A band GO:0031672 | 5.2 | 16 | 3.1 | 2.4e−06 | 3.4e−05 |

**E**

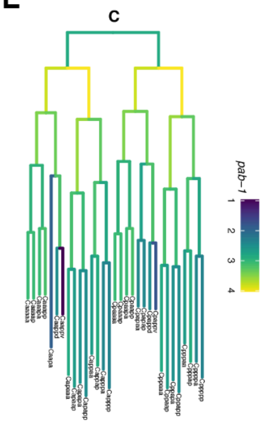


**B**

| Term | Expected | Observed | Enrichment.Fold.Change | P.value | Q.value |
|---|---|---|---|---|---|
| DNA−binding transcription factor activity GO:0003700 | 36.0 | 75 | 2.1 | 5.6e−11 | 1.6e−08 |
| sequence−specific DNA binding GO:0043565 | 43.0 | 85 | 2.0 | 7.7e−11 | 1.6e−08 |
| RNA biosynthetic process GO:0032774 | 85.0 | 140 | 1.6 | 6.4e−10 | 6.1e−08 |
| cellular aromatic compound metabolic process GO:0006725 | 200.0 | 275 | 1.4 | 1.0e−09 | 7.1e−08 |
| heterocycle metabolic process GO:0046483 | 200.0 | 274 | 1.4 | 1.4e−09 | 8.2e−08 |
| organic cyclic compound metabolic process GO:1901360 | 200.0 | 275 | 1.4 | 3.5e−09 | 1.7e−07 |
| transcription regulatory region nucleic acid binding GO:0001067 | 31.0 | 61 | 1.9 | 7.3e−08 | 3.0e−06 |
| double−stranded DNA binding GO:0003690 | 36.0 | 67 | 1.9 | 8.9e−08 | 3.2e−06 |
| condensed chromosome centromeric region GO:0000779 | 7.8 | 22 | 2.8 | 6.4e−07 | 2.0e−05 |
| formation of primary germ layer GO:0001704 | 3.3 | 11 | 3.3 | 2.4e−05 | 6.8e−04 |
| recombinational repair GO:0000725 | 5.1 | 14 | 2.8 | 5.3e−05 | 1.4e−03 |
| metabolic process GO:0008152 | 410.0 | 475 | 1.2 | 1.2e−04 | 2.9e−03 |
| protein−DNA complex GO:0032993 | 51.0 | 74 | 1.5 | 3.1e−04 | 6.8e−03 |
| single−stranded DNA binding GO:0003697 | 3.8 | 10 | 2.6 | 7.2e−04 | 1.5e−02 |
| nucleic acid transport GO:0050657 | 6.9 | 15 | 2.2 | 7.4e−04 | 1.5e−02 |
| mRNA transport GO:0051028 | 5.1 | 12 | 2.4 | 8.5e−04 | 1.5e−02 |
| meiotic cell cycle GO:0051321 | 20.0 | 33 | 1.7 | 1.1e−03 | 1.8e−02 |
| RNA localization GO:0006403 | 7.1 | 15 | 2.1 | 1.2e−03 | 1.9e−02 |
| cell−cell signaling by wnt GO:0198738 | 5.5 | 12 | 2.2 | 1.9e−03 | 2.9e−02 |
| import into nucleus GO:0051170 | 5.2 | 11 | 2.1 | 3.7e−03 | 5.3e−02 |

**F**

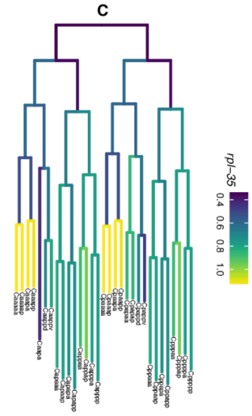


**C**

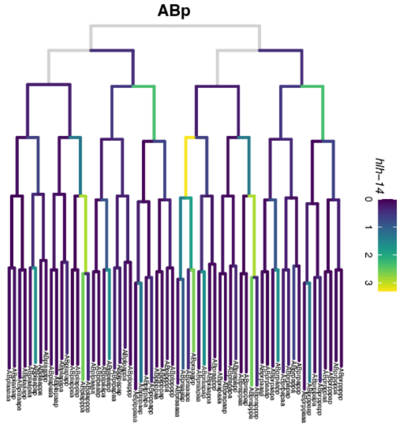


**D**

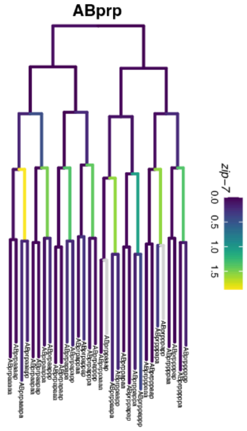


**Supplemental Figure S2. Extended gene ontology analysis results for bulk data and lineage tree examples of highly transient and persistent genes.** (A) Twenty most significantly enriched gene ontology terms for the top 15% stable transcripts. Background set of genes used was all genes that met our moderate mRNA half-life filtering metric. (B) Twenty most significantly enriched gene ontology terms for the top 15% unstable transcripts. Background set of genes used was all genes that met our moderate mRNA half-life filtering metric. (C, D, E, F) Sublineages with coloring representing gene expression from our *C. elegans* embryo single cell atlas (Packer et al. 2019).

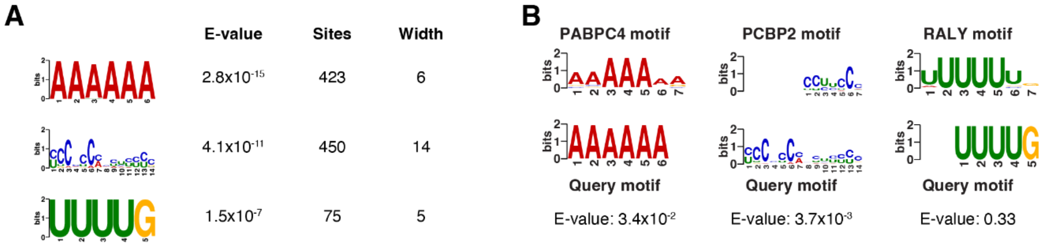


**Supplemental Figure S3. Extended motif analysis results for stable transcripts in the bulk data.** (A) Motifs found to be differentially enriched in the 3’ UTRs of the top 15% stable transcripts using the *de novo* motif-finding program MEME (Bailey et al. 2015), including the E-value, number of sites found, and width for each motif. The 3’ UTRs of the top 15% unstable transcripts were used as control sequences. (B) Mammalian motifs with the highest similarity to the motifs identified in (A) using the Tomtom motif comparison tool against a database of known motifs (Ray et al. 2013).

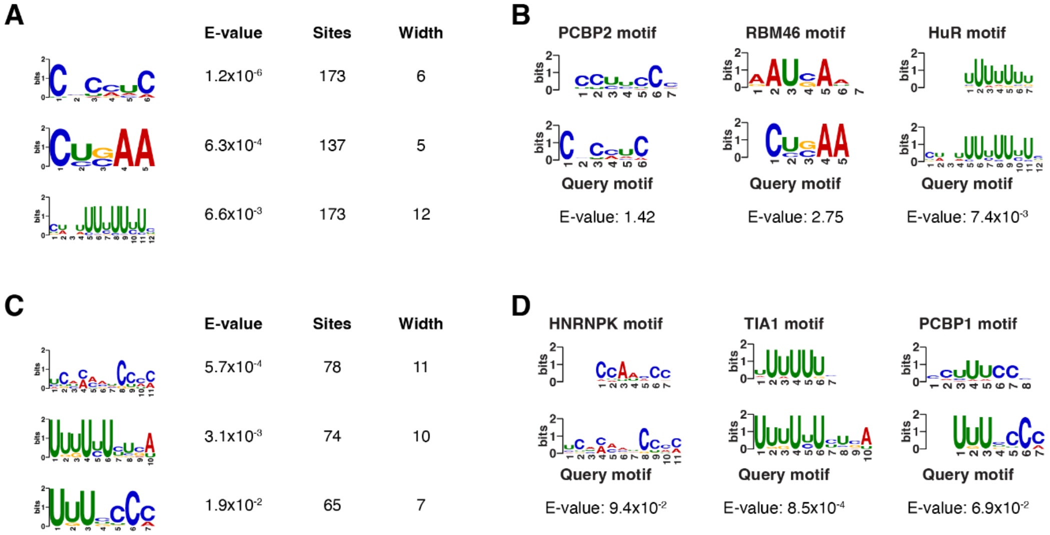


**Supplemental Figure S4. Extended motif analysis results for genes that accumulate to high transcript levels.** (A) Motifs found to be differentially enriched in the 3’ UTRs of genes that accumulate to high transcript levels ~200 minutes past the four-cell stage in a whole embryo RNA-seq dataset (Hashimshony et al. 2015). Motifs were identified using the *de novo* motif-finding program MEME (Bailey et al. 2015). Table includes the E-value, number of sites found, and width for each motif. The 3’ UTRs of genes that accumulate to low transcript levels ~200 minutes were used as control sequences. (B) Mammalian motifs with the highest similarity to the motifs identified in (A) using the Tomtom motif comparison tool against a database of known motifs (Ray et al. 2013). (C) Motifs found to be differentially enriched in the 3’ UTRs of genes that accumulate to high transcript levels ~350 minutes past the four-cell stage in a whole embryo RNA-seq dataset (Hashimshony et al. 2015). Motifs were identified using the *de novo* motif-finding program MEME (Bailey et al. 2015). Table includes the E-value, number of sites found, and width for each motif. The 3’ UTRs of genes that accumulate to low transcript levels ~350 minutes were used as control sequences. (D) Mammalian motifs with the highest similarity to the motifs identified in (C) using the Tomtom motif comparison tool against a database of known motifs (Ray et al. 2013).

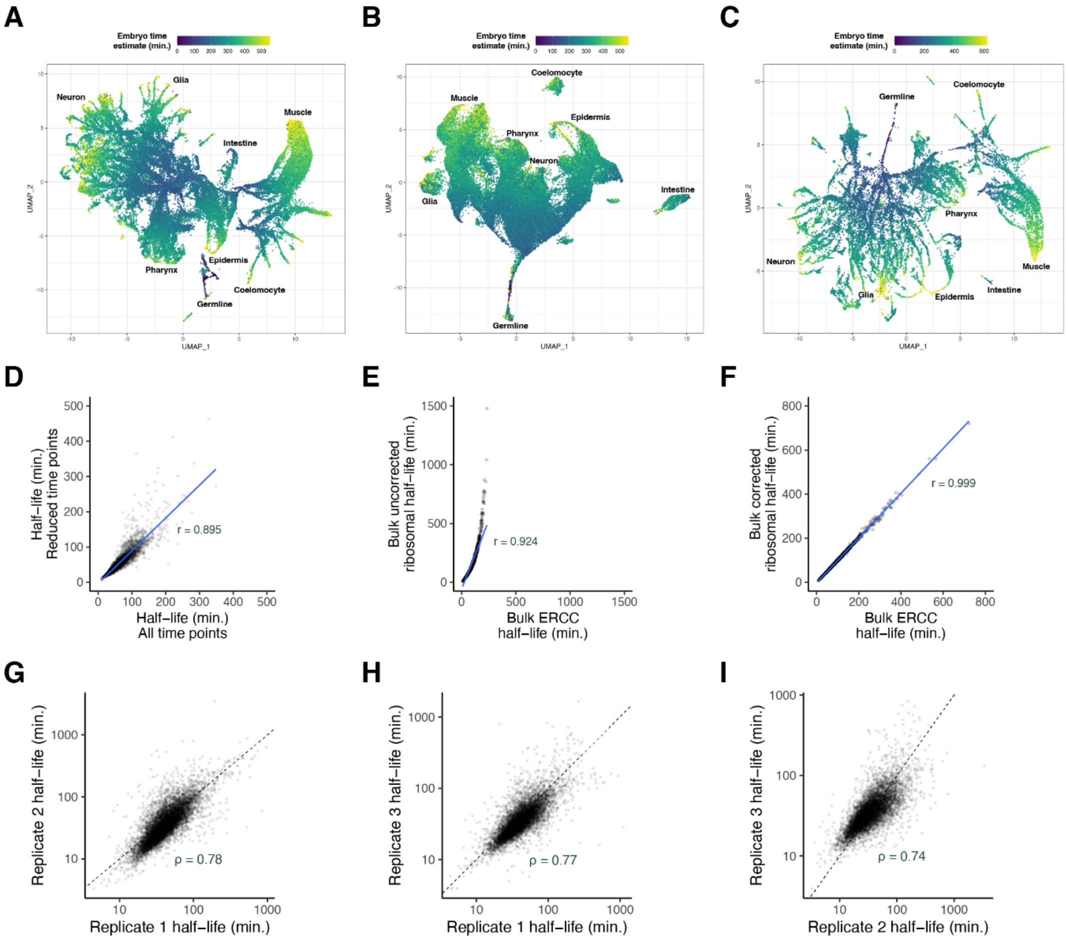


**Supplemental Figure S5. Quality control for transcription inhibition approach paired with single-cell RNA-sequencing.** (A, B, C) Global UMAPs for individual biological replicates, with cells colored by embryo age as estimated from correlations to a whole-embryo RNA-sequencing time series (Hashimshony et al. 2015). Trajectories corresponding to major cell types are labeled. (D) Scatter plot showing the comparison of calculated mRNA half-lives when using all time points (0, 10, 20, 40, 60 minutes) or reduced time points (0, 20, 40 minutes) from the bulk data. Genes were compared if they met the following criteria: coefficient of variation across biological replicates ≤ 50% or the fold-change between the upper limit of their 95% confidence interval and measured half-life ≤ 3. To better include high-stability mRNAs, genes with half-lives > 100 minutes were allowed a looser filtering strategy. Such genes were included if their half-lives had a coefficient of variation ≤ 75% or fold-change between the upper limit of their 95% confidence interval and measured half-life ≤ 4. Pearson's correlation coefficient = 0.895. (E) Scatter plot showing the comparison in calculated mRNA half-lives from the bulk data between gene counts normalized to spike-in ERCC transcripts or transcripts encoding ribosomal proteins. Pearson's correlation coefficient = 0.924. Blue line is the best fit line. (F) Scatter plot showing the comparison in calculated mRNA half-lives from the bulk data between gene counts normalized to spike-in ERCC transcripts or transcripts encoding ribosomal proteins after correcting for their decay. Pearson's correlation coefficient = 0.999. Blue line is the best fit line. (G, H, I) Scatter plots showing the comparison of measured mRNA half-lives between 3 single-cell biological replicates on a log-log scale. Genes were compared if they had UMI > 30 at the 0 minute time point and if their decay fit an exponential decay model $R^2$ ≥ 0.75 for each replicate. Spearman correlation coefficient is displayed for each pairwise comparison. Dashed line is the x = y line.

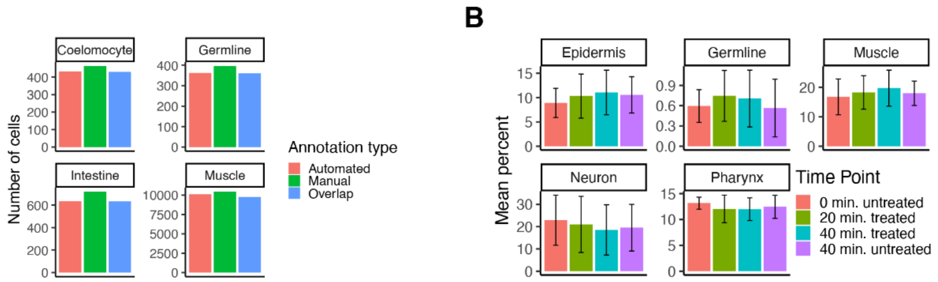


C

| Group1 | Group2 | n1 | n2 | p | p.adjusted |
|---|---|---|---|---|---|
| Muscle | Germline | 4350 | 802 | 2.21e−99 | 1.77e−98 |
| Muscle | Epidermis | 4350 | 3717 | 9.58e−06 | 1.92e−05 |
| Muscle | Neuron | 4350 | 4762 | 7.68e−30 | 3.84e−29 |
| Muscle | Pharynx | 4350 | 4700 | 7.00e−03 | 7.00e−03 |
| Germline | Epidermis | 802 | 3717 | 2.72e−75 | 1.90e−74 |
| Germline | Neuron | 802 | 4762 | 1.36e−186 | 1.36e−185 |
| Germline | Pharynx | 802 | 4700 | 1.78e−118 | 1.60e−117 |
| Epidermis | Neuron | 3717 | 4762 | 2.02e−55 | 1.21e−54 |
| Epidermis | Pharynx | 3717 | 4700 | 6.63e−13 | 1.99e−12 |
| Neuron | Pharynx | 4762 | 4700 | 1.00e−18 | 4.00e−18 |

**Supplemental Figure S6. Quality control for transcription inhibition approach paired with single-cell RNA-sequencing continued.** (A) Bar plot comparing the number of cells from the first biological replicate annotated as coelomocyte, germline, intestine, or muscle based on manual annotation using marker genes or automated annotation using Seurat. (B) Bar plot showing the mean percentage of cells coming from the epidermis, germline, muscle, neuron, and pharynx within each biological replicate, separated by time point. Error bars represent standard deviation between the three biological replicates. (C) Table comparing the mRNA half-life distributions between epidermis, germline, muscle, neuron, and pharynx and whether the distributions are statistically significant from one another. P-values comparing median half-lives were calculated using the Wilcoxon rank sum test.

**A**

| Term | Expected | Observed | Enrichment.Fold.Change | P.value | Q.value |
|---|---|---|---|---|---|
| structural constituent of chromatin GO:0030527 | 1.50 | 10 | 6.7 | 1.2e−07 | 3.5e−05 |
| protein heterodimerization activity GO:0046982 | 2.20 | 10 | 4.6 | 6.9e−06 | 9.8e−04 |
| transmembrane transport GO:0055085 | 9.10 | 20 | 2.2 | 3.3e−04 | 3.2e−02 |
| passive transmembrane transporter activity GO:0022803 | 1.80 | 7 | 3.8 | 3.9e−04 | 3.2e−02 |
| symporter activity GO:0015293 | 0.42 | 3 | 7.1 | 5.0e−04 | 3.2e−02 |
| transporter activity GO:0005215 | 8.80 | 19 | 2.2 | 5.7e−04 | 3.2e−02 |

**B**

| Term | Expected | Observed | Enrichment.Fold.Change | P.value | Q.value |
|---|---|---|---|---|---|
| cilium organization GO:0044782 | 2.400 | 19 | 8.1 | 2.5e−14 | 7.1e−12 |
| non−motile cilium assembly GO:1905515 | 1.100 | 12 | 11.0 | 6.4e−12 | 9.1e−10 |
| ciliary basal body GO:0036064 | 1.100 | 9 | 8.4 | 2.9e−08 | 2.7e−06 |
| cell projection GO:0042995 | 12.000 | 31 | 2.6 | 5.0e−07 | 3.6e−05 |
| cell projection organization GO:0030030 | 10.000 | 27 | 2.6 | 1.4e−06 | 8.0e−05 |
| microtubule−based transport GO:0099111 | 1.800 | 9 | 5.0 | 8.0e−06 | 3.8e−04 |
| non−motile cilium GO:0097730 | 1.200 | 7 | 6.0 | 1.2e−05 | 4.8e−04 |
| ciliary plasm GO:0097014 | 0.690 | 5 | 7.3 | 2.9e−05 | 1.0e−03 |
| taxis GO:0042330 | 5.500 | 15 | 2.7 | 1.3e−04 | 4.0e−03 |
| supramolecular polymer GO:0099081 | 8.100 | 18 | 2.2 | 5.2e−04 | 1.5e−02 |
| monoatomic ion homeostasis GO:0050801 | 2.400 | 8 | 3.3 | 5.5e−04 | 1.5e−02 |
| inorganic ion import across plasma membrane GO:0099587 | 0.490 | 3 | 6.1 | 9.5e−04 | 2.2e−02 |
| neurotransmitter receptor activity involved in regulation of postsynaptic membrane potential GO:0099529 | 0.098 | 1 | 10.0 | 2.4e−03 | 5.2e−02 |
| transmembrane transport GO:0055085 | 10.000 | 19 | 1.9 | 3.4e−03 | 6.9e−02 |
| synaptic signaling GO:0099536 | 3.700 | 9 | 2.4 | 3.9e−03 | 7.3e−02 |
| sodium ion transport GO:0006814 | 0.730 | 3 | 4.1 | 5.1e−03 | 9.0e−02 |
| passive transmembrane transporter activity GO:0022803 | 2.700 | 7 | 2.6 | 5.1e−03 | 9.0e−02 |
| gated channel activity GO:0022836 | 1.200 | 4 | 3.4 | 5.5e−03 | 9.0e−02 |
| monocarboxylic acid biosynthetic process GO:0072330 | 0.780 | 3 | 3.8 | 6.5e−03 | 9.7e−02 |
| chemosensory behavior GO:0007635 | 1.200 | 4 | 3.3 | 6.5e−03 | 9.7e−02 |

**C**

| Term | Expected | Observed | Enrichment.Fold.Change | P.value | Q.value |
|---|---|---|---|---|---|
| structural constituent of chromatin GO:0030527 | 0.59 | 4 | 6.8 | 0.00016 | 0.045 |
| transmembrane transport GO:0055085 | 8.00 | 18 | 2.3 | 0.00039 | 0.055 |
| peptidase inhibitor activity GO:0030414 | 0.43 | 3 | 7.0 | 0.00049 | 0.055 |
| endopeptidase regulator activity GO:0061135 | 0.43 | 3 | 7.0 | 0.00049 | 0.055 |
| extrinsic component of cytoplasmic side of plasma membrane GO:0031234 | 0.48 | 3 | 6.2 | 0.00085 | 0.055 |
| extracellular region GO:0005576 | 1.60 | 6 | 3.7 | 0.00088 | 0.055 |
| proteoglycan metabolic process GO:0006029 | 0.54 | 3 | 5.6 | 0.00140 | 0.055 |
| inorganic ion import across plasma membrane GO:0099587 | 0.27 | 2 | 7.4 | 0.00140 | 0.055 |
| protein serine kinase activity GO:0106310 | 3.40 | 9 | 2.6 | 0.00210 | 0.066 |
| potassium ion transmembrane transport GO:0071805 | 0.32 | 2 | 6.2 | 0.00280 | 0.078 |
| chloride channel complex GO:0034707 | 0.11 | 1 | 9.3 | 0.00290 | 0.078 |
| monoatomic anion transport GO:0006820 | 0.65 | 3 | 4.6 | 0.00290 | 0.078 |
| monoatomic ion homeostasis GO:0050801 | 2.00 | 6 | 2.9 | 0.00380 | 0.082 |
| cytosolic large ribosomal subunit GO:0022625 | 0.70 | 3 | 4.3 | 0.00400 | 0.082 |
| endoplasmic reticulum subcompartment GO:0098827 | 7.00 | 14 | 2.0 | 0.00430 | 0.082 |
| transporter activity GO:0005215 | 7.70 | 15 | 1.9 | 0.00450 | 0.082 |
| symporter activity GO:0015293 | 0.38 | 2 | 5.3 | 0.00460 | 0.082 |
| sodium ion transport GO:0006814 | 0.38 | 2 | 5.3 | 0.00460 | 0.082 |
| nuclear outer membrane−endoplasmic reticulum membrane network GO:0042175 | 7.30 | 14 | 1.9 | 0.00620 | 0.092 |

**Supplemental Figure S7. Extended gene ontology analysis results for genes with more rapid mRNA decay over time.** (A) Significantly enriched gene ontology terms for the top 5% of genes with faster decay in Middle-stage cells compared to Early-stage cells. Background set of genes used was shared genes between Early- and Middle-stage cells that met our moderate mRNA half-life filtering metric. (B) Twenty most significantly enriched gene ontology terms for the top 5% of genes with faster decay in Late-stage cells compared to Middle-stage cells. Background set of genes used was shared genes between Middle- and Late-stage cells that met our moderate mRNA half-life filtering metric. (C) Significantly enriched gene ontology terms for the top 5% of genes with faster decay in Late-stage cells compared to Early-stage cells. Background set of genes used was shared genes between Early- and Late-stage cells that met our moderate mRNA half-life filtering metric.

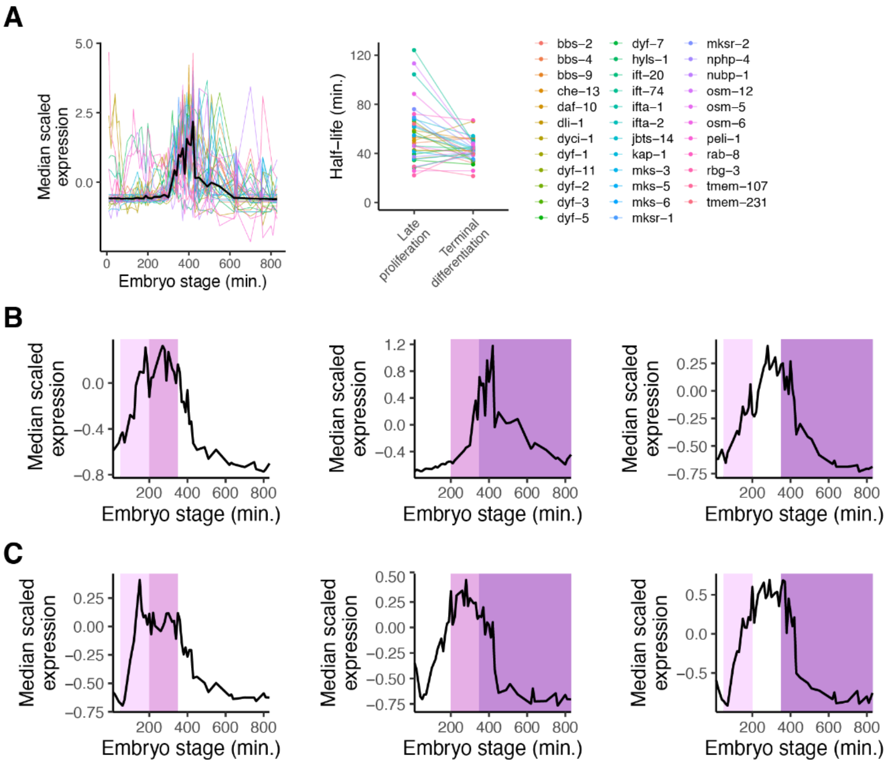


**Supplemental Figure S8. Extended analyses for genes with differential mRNA decay over time.** (A) *Left*. Median scaled expression of core cilia component genes using data from a whole-embryo RNA-sequencing time series (Hashimshony et al. 2015). *Right*. Plot displaying the change in mRNA half-lives from Middle to Late stage for core cilia component genes. (B) Median scaled expression of zygotic-only genes in the top 5% of genes with faster mRNA decay in a later stage compared to in an earlier stage. Pink shading spans the Early stage, light purple shading spans the Middle stage, and dark purple shading spans the Late stage. (C) Median scaled expression of zygotic-only genes in the top 5% of genes with slower mRNA decay in a later stage compared to in an earlier stage. Pink shading spans the Early stage, light purple shading spans the Middle stage, and dark purple shading spans the Late stage.

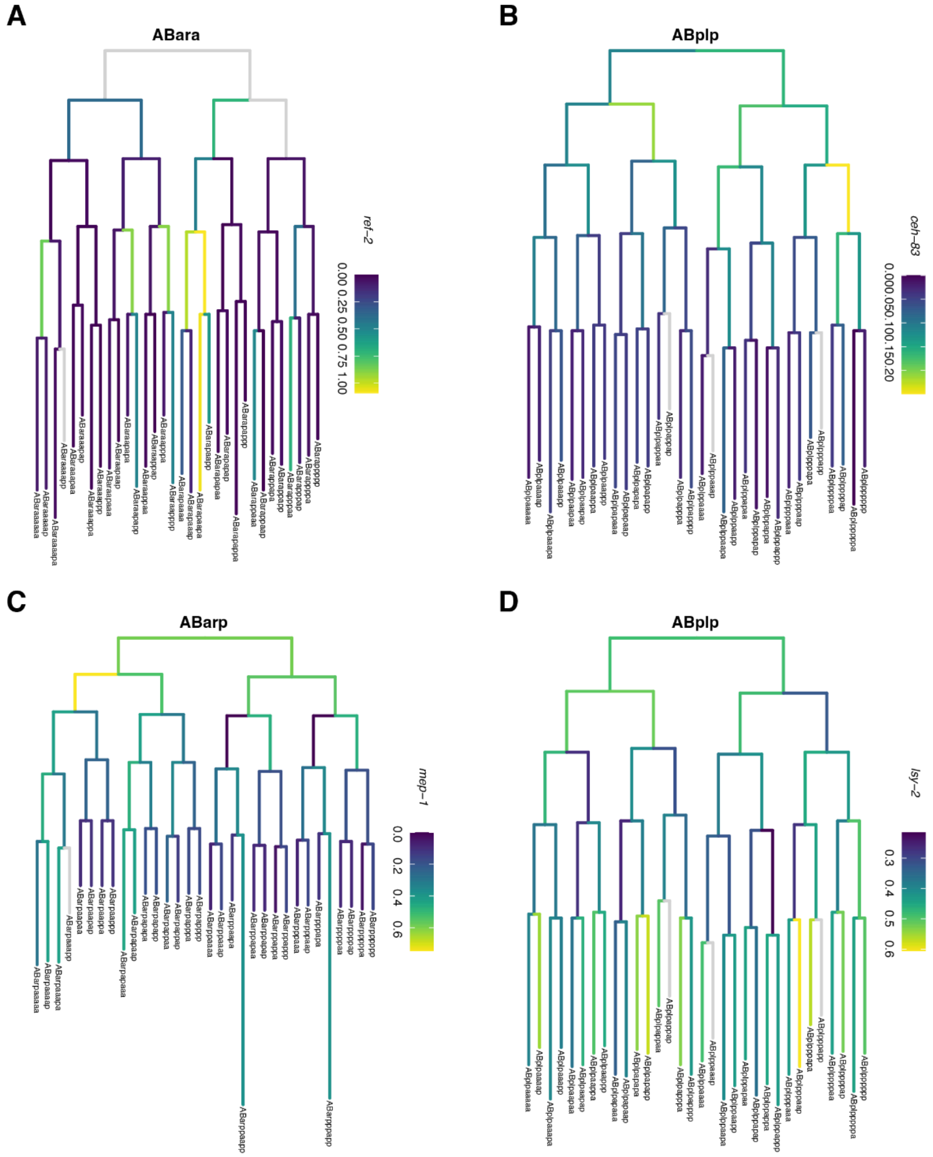


**Supplemental Figure S9. Lineage tree examples of transcription factor genes with transient or persistent mRNA expression.** (A) Lineage tree for the ABara sublineage with coloring representing *ref-2* mRNA expression from our *C. elegans* embryo single cell atlas (Packer et al. 2019). (B) Lineage tree for the ABplp sublineage with coloring representing *ceh-83* mRNA expression from our *C. elegans* embryo single cell atlas (Packer et al. 2019). (C) Lineage tree for the ABarp sublineage with coloring representing *mep-1* mRNA expression from our *C. elegans* embryo single cell atlas (Packer et al. 2019). (D) Lineage tree for the ABplp sublineage with coloring representing *lsy-2* mRNA expression from our *C. elegans* embryo single cell atlas (Packer et al. 2019).

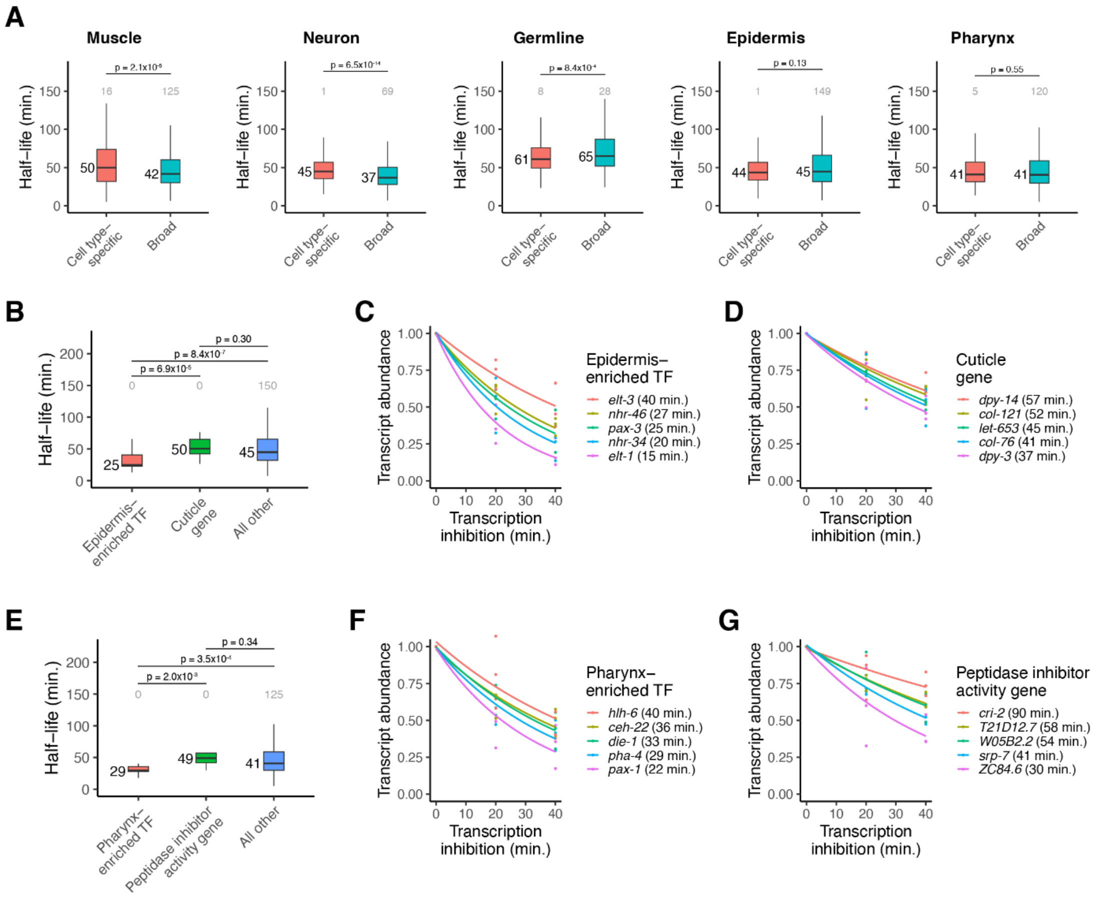


**Supplemental Figure S10. mRNA half-lives of cell type-specific genes.** (A) Box plots showing the mRNA half-life distributions of cell type-specific and broadly expressed genes within muscle, germline, epidermis, neuron, and pharynx cells. (B) Box plots showing the epidermis-specific mRNA half-life distributions of epidermis-enriched transcription factor genes, cuticle genes, and all other genes. (C) Scatter plot of the normalized transcript abundance of the epidermis-enriched transcription factor genes *elt-3*, *nhr-46*, *pax-3*, *nhr-34*, *elt-1* throughout a 40 minute transcription inhibition time course in epidermal cells. Each point represents normalized transcript abundance from one of three biological replicates. (D) Scatter plot of the normalized transcript abundance of the cuticle genes *dpy-14*, *col-121*, *let-653*, *col-76*, *dpy-3* throughout a 40 minute transcription inhibition time course in epidermal cells. Each point represents normalized transcript abundance from one of three biological replicates. (E) Box plots showing the pharynx-specific mRNA half-life distributions of pharynx-enriched transcription factor genes, peptidase inhibitor activity genes, and all other genes. (F) Scatter plot of the normalized transcript abundance of the pharynx-enriched transcription factor genes *hlh-6*, *ceh-22*, *die-1*, *pha-4, pax-1* throughout a 40 minute transcription inhibition time course in pharynx cells. Each point represents normalized transcript abundance from one of three biological replicates. (G) Scatter plot of the normalized transcript abundance of the peptidase inhibitor activity genes *cri-2*, *T21D12.7*, *W05B2.2*, *srp-7*, *ZC84.6* throughout a 40 minute transcription inhibition time course in pharynx cells. Each point represents normalized transcript abundance from one of three biological replicates. Numbers to the left of the box plots are median half-lives within each group. Numbers above box plots are the number of genes with half-lives greater than 150 minutes within each group. P-values comparing median half-lives were calculated using the Wilcoxon rank sum test.

**A**

| Term | Expected | Observed | Enrichment.Fold.Change | P.value | Q.value |
|---|---|---|---|---|---|
| supramolecular polymer GO:0099081 | 9.9 | 50 | 5.1 | 3.8e−23 | 5.4e−21 |
| striated muscle dense body GO:0055120 | 4.5 | 33 | 7.4 | 3.7e−22 | 3.5e−20 |
| myofibril GO:0030016 | 3.6 | 29 | 8.1 | 2.8e−21 | 2.0e−19 |
| extracellular region GO:0005576 | 3.6 | 25 | 7.0 | 1.1e−16 | 6.4e−15 |
| A band GO:0031672 | 1.7 | 17 | 10.0 | 6.0e−16 | 2.8e−14 |
| muscle system process GO:0003012 | 1.7 | 17 | 9.9 | 1.5e−15 | 6.0e−14 |
| sarcomere organization GO:0045214 | 1.5 | 16 | 10.0 | 2.6e−15 | 9.1e−14 |
| gated channel activity GO:0022836 | 2.3 | 18 | 8.0 | 5.6e−14 | 1.8e−12 |
| passive transmembrane transporter activity GO:0022803 | 4.2 | 24 | 5.7 | 1.4e−13 | 3.9e−12 |
| muscle cell development GO:0055001 | 2.1 | 17 | 7.9 | 2.6e−13 | 6.7e−12 |

**B**

| Term | Expected | Observed | Enrichment.Fold.Change | P.value | Q.value |
|---|---|---|---|---|---|
| cell projection organization GO:0030030 | 14.0 | 78 | 5.4 | 4.7e−38 | 3.4e−36 |
| cilium organization GO:0044782 | 3.8 | 41 | 11.0 | 4.8e−38 | 3.4e−36 |
| cell projection GO:0042995 | 18.0 | 86 | 4.8 | 9.8e−37 | 4.7e−35 |
| ciliary basal body GO:0036064 | 2.0 | 25 | 13.0 | 1.3e−27 | 5.3e−26 |
| non−motile cilium assembly GO:1905515 | 1.7 | 21 | 12.0 | 3.9e−23 | 1.4e−21 |
| ciliary plasm GO:0097014 | 1.1 | 15 | 14.0 | 2.2e−19 | 6.8e−18 |
| taxis GO:0042330 | 8.1 | 37 | 4.6 | 2.2e−16 | 6.1e−15 |
| neuron development GO:0048666 | 8.8 | 37 | 4.2 | 5.8e−15 | 1.5e−13 |
| microtubule−based transport GO:0099111 | 2.6 | 19 | 7.3 | 4.6e−14 | 1.1e−12 |
| extracellular region GO:0005576 | 3.2 | 17 | 5.3 | 6.6e−10 | 1.4e−08 |

**C**

| Term | Expected | Observed | Enrichment.Fold.Change | P.value | Q.value |
|---|---|---|---|---|---|
| extracellular region GO:0005576 | 3.70 | 24 | 6.6 | 1.5e−15 | 8.4e−14 |
| molting cycle GO:0042303 | 2.50 | 19 | 7.7 | 1.9e−14 | 8.8e−13 |
| organic acid metabolic process GO:0006082 | 9.40 | 37 | 3.9 | 6.0e−14 | 2.4e−12 |
| structural constituent of cuticle GO:0042302 | 0.72 | 9 | 13.0 | 1.4e−11 | 5.0e−10 |
| oxidoreductase activity acting on CH−OH group of donors GO:0016614 | 1.90 | 11 | 5.8 | 1.0e−07 | 3.3e−06 |
| DNA−binding transcription factor activity GO:0003700 | 9.30 | 27 | 2.9 | 1.4e−07 | 4.1e−06 |
| microbody GO:0042579 | 1.60 | 10 | 6.1 | 1.7e−07 | 4.3e−06 |
| zinc ion binding GO:0008270 | 7.90 | 24 | 3.0 | 2.3e−07 | 5.3e−06 |
| iron ion binding GO:0005506 | 1.20 | 8 | 6.5 | 1.1e−06 | 2.3e−05 |
| peptidase inhibitor activity GO:0030414 | 0.59 | 5 | 8.5 | 5.4e−06 | 1.1e−04 |

**D**

| Term | Expected | Observed | Enrichment.Fold.Change | P.value | Q.value |
|---|---|---|---|---|---|
| serine−type endopeptidase inhibitor activity GO:0004867 | 0.18 | 7 | 40.0 | 5.4e−14 | 7.7e−12 |
| peptidase inhibitor activity GO:0030414 | 0.26 | 7 | 26.0 | 2.5e−11 | 2.4e−09 |
| endopeptidase regulator activity GO:0061135 | 0.29 | 7 | 24.0 | 6.3e−11 | 4.5e−09 |
| DNA−binding transcription factor activity GO:0003700 | 4.10 | 19 | 4.6 | 6.6e−09 | 3.8e−07 |
| transcription regulatory region nucleic acid binding GO:0001067 | 3.80 | 17 | 4.5 | 5.7e−08 | 2.7e−06 |
| sequence−specific DNA binding GO:0043565 | 5.20 | 19 | 3.7 | 3.1e−07 | 1.2e−05 |
| double−stranded DNA binding GO:0003690 | 4.40 | 17 | 3.9 | 5.6e−07 | 2.0e−05 |
| pharynx development GO:0060465 | 0.53 | 6 | 11.0 | 6.2e−07 | 2.0e−05 |
| cell surface GO:0009986 | 0.46 | 5 | 11.0 | 4.6e−06 | 1.3e−04 |
| external encapsulating structure GO:0030312 | 0.46 | 4 | 8.6 | 7.8e−05 | 2.0e−03 |

**E**

| Term | Expected | Observed | Enrichment.Fold.Change | P.value | Q.value |
|---|---|---|---|---|---|
| mRNA 3'−UTR binding GO:0003730 | 4.3 | 8 | 1.8 | 0.0014 | 0.014 |
| membrane−enclosed lumen GO:0031974 | 73.0 | 90 | 1.2 | 0.0019 | 0.018 |
| single−stranded DNA binding GO:0003697 | 3.4 | 6 | 1.8 | 0.0061 | 0.056 |
| recombinational repair GO:0000725 | 3.4 | 6 | 1.8 | 0.0061 | 0.056 |
| cell part morphogenesis GO:0032990 | 3.4 | 6 | 1.8 | 0.0061 | 0.056 |
| neuron development GO:0048666 | 2.9 | 5 | 1.7 | 0.0130 | 0.110 |
| identical protein binding GO:0042802 | 9.7 | 14 | 1.4 | 0.0140 | 0.110 |
| import into nucleus GO:0051170 | 5.3 | 8 | 1.5 | 0.0250 | 0.200 |
| ATP−dependent activity acting on RNA GO:0008186 | 5.3 | 8 | 1.5 | 0.0250 | 0.200 |
| extracellular region GO:0005576 | 2.4 | 4 | 1.7 | 0.0260 | 0.200 |

**Supplemental Figure S11. Gene ontology analysis of cell type-specific genes.** (A) Ten most significantly enriched gene ontology terms for muscle-specific genes that met our moderate mRNA half-life filtering metric within muscle cells. Background set of genes used was all genes that met our moderate mRNA half-life filtering metric within muscle cells. (B) Ten most significantly enriched gene ontology terms for neuron-specific genes that met our moderate mRNA half-life filtering metric within neuronal cells. Background set of genes used was all genes that met our moderate mRNA half-life filtering metric within neuron cells. (C) Ten most significantly enriched gene ontology terms for epidermis-specific genes that met our moderate mRNA half-life filtering metric within epidermal cells. Background set of genes used was all genes that met our moderate mRNA half-life filtering metric within epidermal cells. (D) Ten most significantly enriched gene ontology terms for pharynx-specific genes that met our moderate mRNA half-life filtering metric within pharynx cells. Background set of genes used was all genes that met our moderate mRNA half-life filtering metric within pharynx cells. (E) Ten most significantly enriched gene ontology terms for germline-specific genes that met our moderate mRNA half-life filtering metric within germline cells. Background set of genes used was all genes that met our moderate mRNA half-life filtering metric within germline cells.
